# HIV-1 Assembly is Assisted by IRSp53 Through an Interaction that Extends Gag Conformation

**DOI:** 10.64898/2026.10.05.756744

**Authors:** Curt Waltmann, Peggy Merida, Puja Banjeree, Karolina Buresova, Chloé Poussineau, Julien Burlaud-Gaillard, Raviteja Gurram, Jeriann Beiter, Cyril Favard, Delphine Muriaux, Gregory A. Voth

## Abstract

Budding of the immature virion is an essential part of the HIV-1 life cycle. However, understanding and inhibiting this process has been challenging, as both simulations and theoretical studies suggest that the energy required to bend the membrane is too high for the complete assembly of a spherical viral particle at the plasma membrane without the assistance of cellular host factors. Recent results suggest that the I-BAR protein IRSp53 may be one such host factor, but the molecular mechanism by which the Gag-IRSp53 interaction promotes HIV-1 assembly and budding remains poorly understood. We employ molecular dynamics simulations, starting from AlphaFold models, to show that the SH3 domain of IRSp53 interacts directly with the MACANTD domain of Gag and we experimentally validate this interaction. Our simulations show that the specificity of the interaction with IRSp53 arises from its ability to extend the MACANTD Gag conformation in a manner similar to RNA binding, which is known to activate Gag assembly. We then computationally design HIV-1 Gag mutants that, as confirmed by simulations, do not undergo conformational changes upon interaction with theIRSp53 SH3 domain. Experimentally, these Gag mutants are tested and shown to accumulate in cells. In addition, for these mutants we observe large quantities of Gag assembling at the plasma membrane but exhibiting large budding defects. These results validate the proposed mechanism by which the Gag–IRSp53 interaction promotes completion of HIV-1 budding and suggest new routes for inhibiting viral assembly or budding.

## Introduction

Assembly of the viral Gag polyprotein is a crucial step in the HIV-1 life cycle(*1*). Gag proteins assemble at the cell membrane forming a hexameric, albeit defective(*2*), lattice due to interactions between Gag molecules in the MA, CANTD, and SP2 domains of the polyprotein(*3, 4*). This lattice forms the spherical, immature particle that packages the viral RNA(*5, 6*) and buds from the cell membrane to infect other cells and continue the viral life cycle(*7*). However, recent studies suggest that the viral RNA also changes the conformation of the Gag molecules themselves, from a compact to extended state, and this facilitates Gag assembly(*8, 9*). While the role of RNA in accelerating Gag assembly is understood, how the Gag lattice generates sufficient curvature to bud from the membrane is not(*10*). Previous coarse-grained (CG) molecular dynamics (MD) simulations of Gag assembly, which included an RNA model, showed stalled budding at the plasma membrane(*11*). Other theoretical studies(*12*) agree that the Gag lattice alone cannot curve the membrane sufficiently to form the viral particle. This is also the case for membrane scission, where Gag “hijacks” the ESCRT machinery (*13, 14*) to complete membrane scission and it seems Gag uses a similar approach for membrane bending. Our previous work(*15*) showed that knocking out the membrane-bending I-BAR protein IRSp53(*16–20*) stalled HIV-1 particle formation, suggesting that this host protein helps the virus to curve the membrane. Strangely, another closely related I-BAR protein(*21*), IRTKS, does not have the same effect on budding even though both proteins associate with Gag in immunoprecipitation experiments and are thought to be functionally redundant(*22*).

Thus, a number of questions remain in relation to the role of IRSp53 in HIV-1 particle formation. For example, what is the nature of the interaction between Gag and IRSp53? Is it membrane-mediated? And, if so, is it related to the clustering of phosphatidylinositol 4,5-bisphosphate (PI(4,5)P2) lipids (*23–25*)? Or is it a direct protein-protein interaction? If the interaction is direct, which of the many domains of these 2 polyproteins is involved? Why does IRTKS not show the same behavior? Aside from a greater understanding of IRSp53 and Gag, answering these questions may also lead to the discovery of antiviral compounds targeting HIV assembly and budding and increase our understanding of the general mechanisms by which viruses evolve to use host cell factors. Moreover, can we utilize computational methods to understand the molecular mechanisms and predict mutations that can experimentally confirm different hypotheses?

Computational methods are widely used in bioscience today. In the past 20 years, Nobel prizes have been given for the development of two major techniques: all-atom MD(*26*), which allows one to sample the many different conformational states of a system, and inverse folding models like AlphaFold(*27–30*), which utilize protein structure data to predict the folded structures of proteins given a primary sequence. These methods are complementary as all-atom MD can accurately sample many conformations but requires an accurate initial state to begin with and is computationally intensive(*31*). Coarse-grained (CG) MD can be used to increase the length and time scales of the simulation at the possible expense of reduced accuracy(*32*). AlphaFold and related models can quickly predict possible structures, but generally not a whole conformational ensemble. Artificial Intelligence (AI) inspired techniques such as diffusion models are being developed to remedy this(*33–37*) but are not yet as accurate as MD at generating ensembles. However, by combining structural predictions with all-atom MD, one can develop a workflow in which AlphaFold predicts initial binding states that are then simulated with MD(*38*) to confirm strong interactions, a workflow that has recently been shown to correct issues with AlphaFold structures with short simulation times(*39*). The strongly interacting state(s) can then be run for much longer MD time periods to sample the entire ensemble of conformations produced on nano to microsecond time scales.

In this work, we use such a workflow to computationally predict a direct interaction between the IRSp53 SH3 domain and the Gag MACANTD. This interaction is then verified in cells through immunoprecipitation experiments. Further simulation shows that this interaction changes the conformational states of Gag, making the protein more extended rather than compact. Importantly, this change does not occur when Gag is bound to the SH3 domain of IRTKS, which was previously established as an experimental control. Instead, the Gag conformations are like the extended conformations observed when Gag interacts with RNA, which is known to accelerate Gag assembly. Analysis of the simulations allows us to predict mutations to Gag which will not eliminate the interaction with IRSp53 but alter it such that the Gag conformation is no longer similar to the RNA bound activated state and more like the IRTKS bound state. We confirm these effects in silico by mutating Gag and observing the expected compact conformations. Finally, we experimentally verify the effect of these mutations using TIRF-M and TEM imaging of altered Gag budding. These results suggest that our mutations have indeed altered the Gag assembly and budding process and thus IRSp53 facilitates wild-type (WT) Gag assembly by affecting its conformational state.

## Results

### The full length protein IRSp53 is needed to interact with HIV-1 Gag

First, we confirmed that our system of transfected eukaryotic cells efficiently produces HIV-1 Gag virus-like particles (VLPs) and that Gag expression promotes the release of IRSp53 in association with these VLPs (Figure 1A) as it was reported before(*15*). Furthermore, we confirmed that IRSp53 localizes to HIV-1 Gag budding sites in these cells (Figure 1B). Confocal and STED microscopy showed that endogenous IRSp53, detected by immunostaining, was localized at or at the vicinity of HIV-1 Gag budding sites (detected by Gag(i)GFP), consistent with previous reports (Inamdar et al. 2021(*15*)) or either in F-actin enriched membranes (as reported in Scita 2008(*40*)).

**Figure 1:**
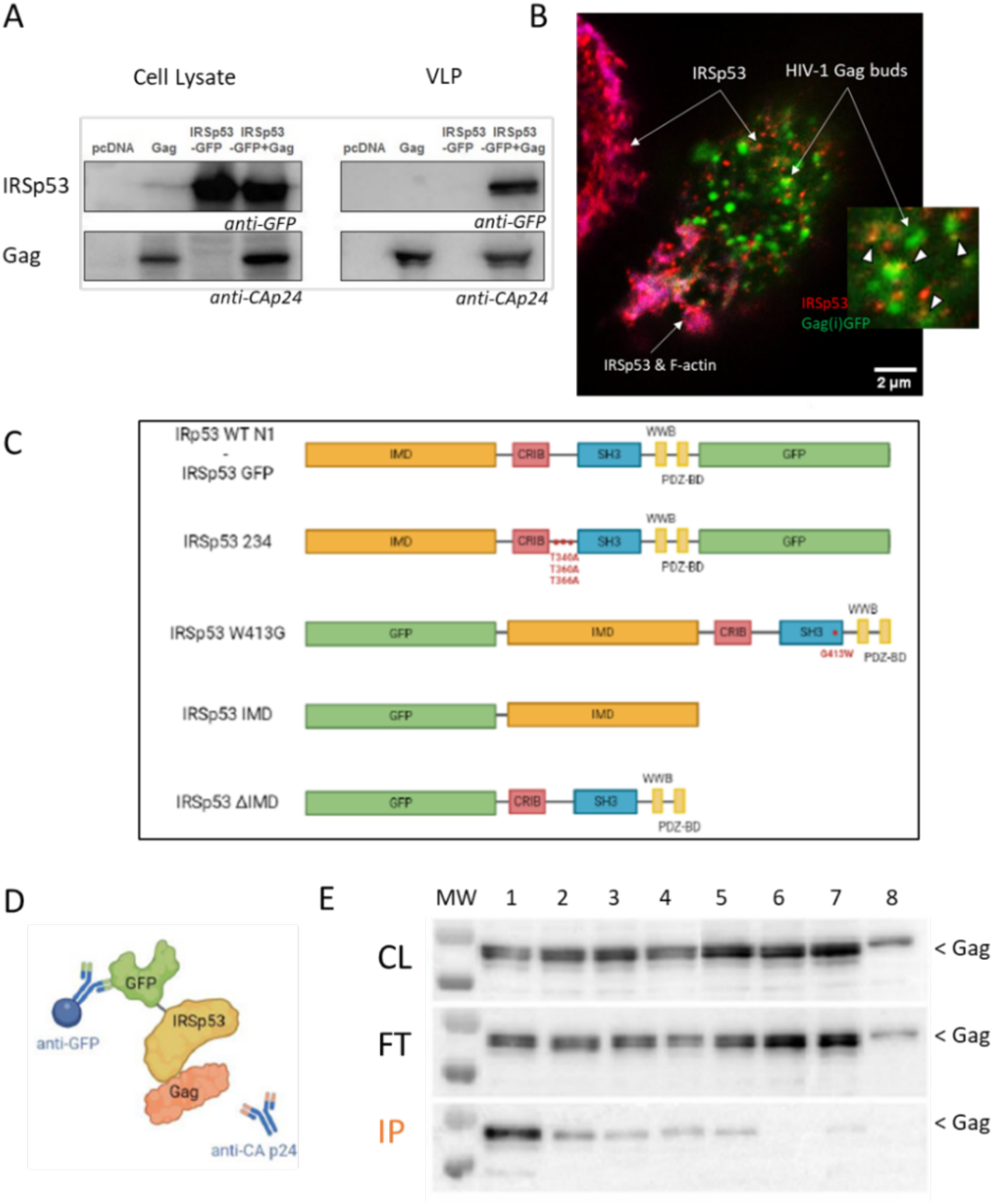
Full length IRSp53 is necessary to interact with HIV-1 Gag. (A) Immunoblots showing Gag VLP production in the absence and in the presence of IIRSp53-GFP. IRSp53-GFP in cell lysates and in VLPs is revealed using anti-GFP and HIV-1 Gag using anti-CAp24 antibodies, as indicated. Results show that IRSp53-GFP is released with Gag VLPs. (B) STED confocal images showing the top view of an HeLa Cells producing HIV-1 Gag(i)GFP and immune-stained for IRSp53 (revealed using anti-IRSp53 and anti-StarRed antibodies). Cells are also labelled with phalloidin-StarOrange for labelling F-actin. A zoomed image shows that IRSp53 is located at close proximity of HIV-1 budding sites, as it was reported in transfected 293THEK cells and Jurkat T cells (Inamdar et al., eLife 2021). STED confocal imaging of HeLa cells (top view) expressing HIV-1 Gag showing HIV-1 Gag buds at the cell surface (green dots) with endogenous IRSp53 (red, anti-IRSp53 labelled with StarRed secondary antibody) and F-actin (Magenta, labelled with Phalloidin). (C) Scheme of the IRSp53-GFP constructs used in this study with the corresponding IRSp53 mutants: Superactive IRSp53-234, W413G, only the N-terminal membrane binding domain of IRSp53, IMD or the C-ter domain dIMD, binding cellular and actin-related factors. (D) Scheme of the GFP-trap system allowing to immunoprecipitated the IRSp53-GFP proteins, WT and mutants, and to reveal by Western blots for their interactions, directly or indirectly, with HIV-1 Gag. (E) Western blots (anti-CAp24) showing that only IRSp53-GFP WT is able to interact well with Gag. Input is the whole Gag present in the cell lysates. FT is the flow-through after application of the cell lysates on the beads. IP is the Gag protein remaining in interaction with the beads carrying IRSp53-GFP or mutants. Lanes 1: WT IRSp53-GFP + Gag; 2: 234+Gag; 3: IMD+Gag; dIMD+Gag; 5:W413G+Gag; 6: GFP+Gag; 7: mock+Gag; 8:none (showing that the upper band is not Gag).

Then, to monitor the interaction between IRSp53-GFP WT or some mutants (Figure 1C) and HIV-1 Gag, we used Immunoprecipitation assays (Figure 1). Briefly, the retroviral Gag and the IRSp53-GFP proteins are overexpressed in eukaryotic cells, lysed and thanks to a GFP trap kit (see Material and Methods section), IRSp53-GFP is trapped on beads thanks to an antibody against the GFP (Figure 1D). After several washes, the beads are run on SDS-PAGE acrylamide gels, and the presence of the Gag protein is revealed by western blot. Following this protocol, we investigated several IRSp53-GFP mutants and if the WT Gag protein was retaining its capacity to IP with IRSp53 (Figure 1E). Lane 1 shows the interaction, directly or indirectly, between HIV-1 Gag and WT IRSp53-GFP. Controls (lanes 6-8) show no interaction with Gag. Any mutation or domain deletion in IRSp53-GFP (lanes 2-5) drastically decreases the interaction with Gag, preventing us from determining any preferential interacting domain of Gag with IRSp53.

We saw that the interaction between Gag and IRSp53 is dependent on the full length of IRSp53 and certainly WT conformation. Since only WT IRSp53 interacts well with HIV-1 Gag, we decided to use molecular modelling to unravel if we can identify any functional interface between Gag and IRSp53.

### Molecular Dynamics Stabilizes the AlphaFold Predicted Interface between HIV-1 Gag CANTD and IRSp53 SH3 Domain

We next sought to determine whether there was a protein-protein interface between HIV-1 Gag and IRSp53. Figure 1 showed that the full length IRSp53 is important for Gag interaction and thus we used the full sequence. We hypothesized that there may be a protein-protein interaction given previous experiments showing their colocalization on a membrane; however, this localization could also be driven by preferences for similar curvatures or lipids like PI(4,5)P2, which is known to localize both proteins. We used structural predictions from the AlphaFold web server on the entire sequences of the HIV-1 Gag and IBAR IRSp53 which have a total length of over 1000 residues. All AlphaFold predictions can be seen in the Supporting Information (SI) Figure S1. highest confidence prediction (see Figure 2A and B) showed, among many other interactions, an interface between the IRSp53 SH3 domain and the HIV-1 CANTD domain. This interaction involves a polyproline helix, which is known to bind SH3 domains(*41*) including the SH3 domain of IRTKS(*42*) (see Figure 2D), and this result was convincing because of the structural similarity with the interaction between the Gag CANTD(*43*) and another host cell factor, CypA, which binds the CANTD later in the HIV-1 lifecycle(*44*) (see Figure 2C).

**Figure 2:**
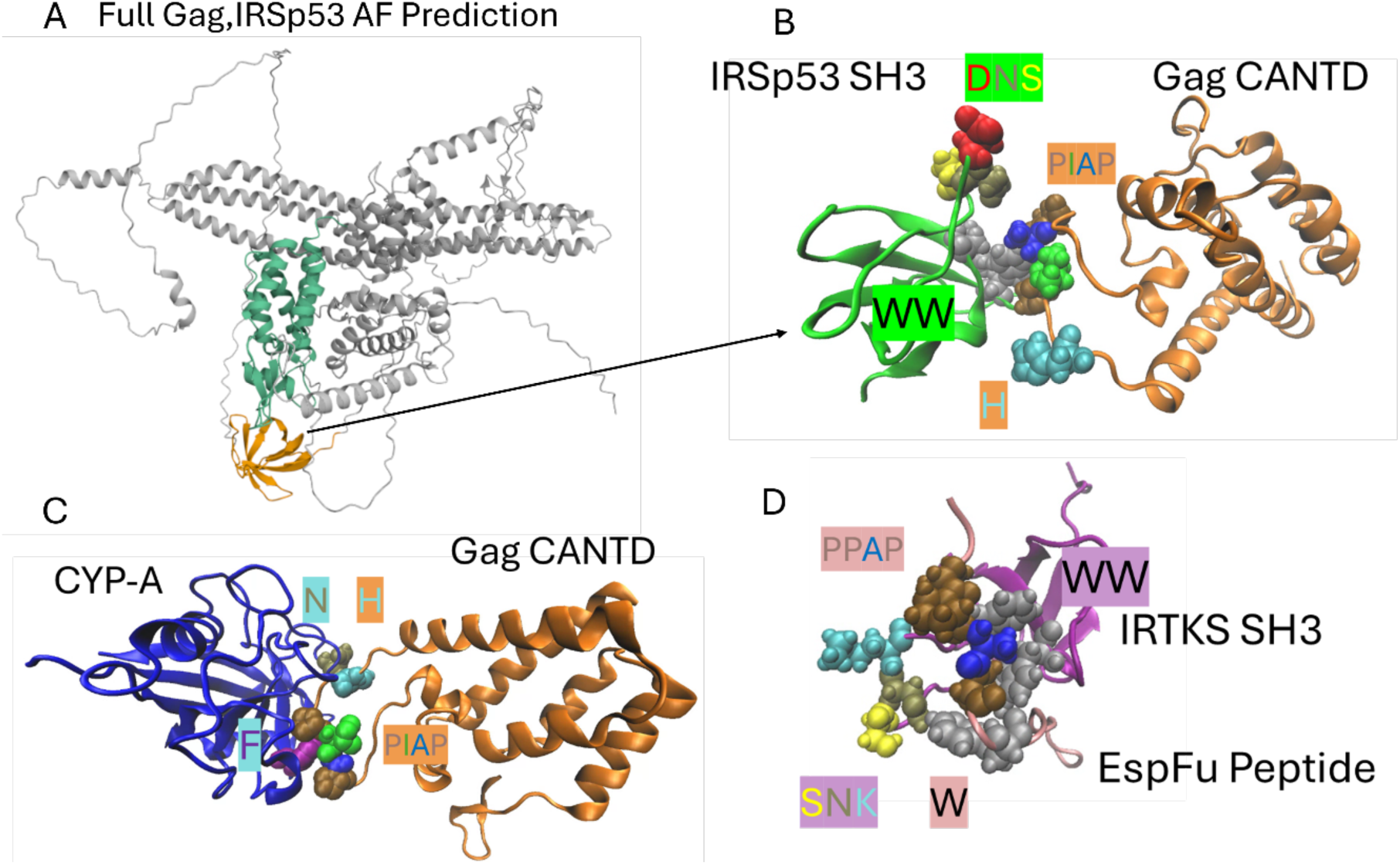
AlphaFold prediction of SH3 domain with Gag CANTD and comparison to previously solved structures. (A) Full structure of the AlphaFold prediction chosen based solely on the SH3 interactions in (B). (B) The structure of the SH3 domain with the Gag CANTD shows a tryptophan in a polyproline helix (PIAP) binding pocket. DNS and H motifs are not interacting although they do after molecular dynamics simulations. (C) Structure of the CypA-Gag interface (PDB:1AK4(43)) shows structural similarity with (B). (D) Structure of the IRTKS SH3 domain with a polyproline helix peptide (PDB: 2KXC(42)). This also has significant structural similarity with (B).

We then set out to confirm the strength of this proposed interaction using all-atom MD. We were somewhat concerned that this interaction only involved the polyproline domain as these are thought to be weak interactions(*45*) that require more residues to be stable as can be seen in the isomorphic solved structures Figure 2C and D. To test this, we fit the known structures of the IRSp53 and IRTKS SH3 domains (from PDBs 3RNJ and 2KXC respectively) and Gag CANTD(*4*) to the structure predicted by AlphaFold. We kept only the Gag CANTD and IRSp53 SH3 domains to make the simulations more computationally efficient. We then ran 6 replicas of MD for 160ns (1 microsecond in total) starting from this same configuration for IRSp53 (Figure 3A) and IRTKS (Figure 3D). In Figure 3B, it is shown that in 4 of the 6 replicas the interface between the IRSp53 and Gag CANTD was stable as quantified by the distance between the initially uninserted TRP59 and Ile222 residues, which decreased due to the need for rearrangement to stabilize the interface. (Incidentally, the residues of the Gag are always numbered by their position in the full sequence shown in the SI figure S4, while the residues in SH3 domains are numbered by their positions within the SH3 domain, also shown in the SI Figure S4.) Figure 3C shows that in each of these stable replicas the DNS motif of IRSp53 SH3 (ASP22, ASN23, SER24), which was not involved in the AlphaFold-predicted interface, begins to interact with some part of the PIAP polyproline helix (Pro221, Ile222, Ala223, Pro224) or His218 shown in Figure 3A. We also see involvement of both tryptophan residues (TRP46 and TRP59) in Figure 3A which was also not predicted by AlphaFold. Together, the two tryptophan’s form a “prong” which can “grab” either of the prolines in the PIAP domain of Gag. These different states are shown by the different distances between W59 and I222 (I of PIAP) in Figure 2B. The same was done for the IRTKS and we observed fewer stable replicas; however, the rearrangement of the interface for those stable replicas was almost identical (see Figure 3D-F). IRTKS has an SNK sequence instead of a DNS sequence, but the interactions were not noticeably different. These additional interactions explain why the interface is stable in a manner consistent with our initial hypothesis that polyproline interactions alone would be insufficient to stabilize the interface.

**Figure 3:**
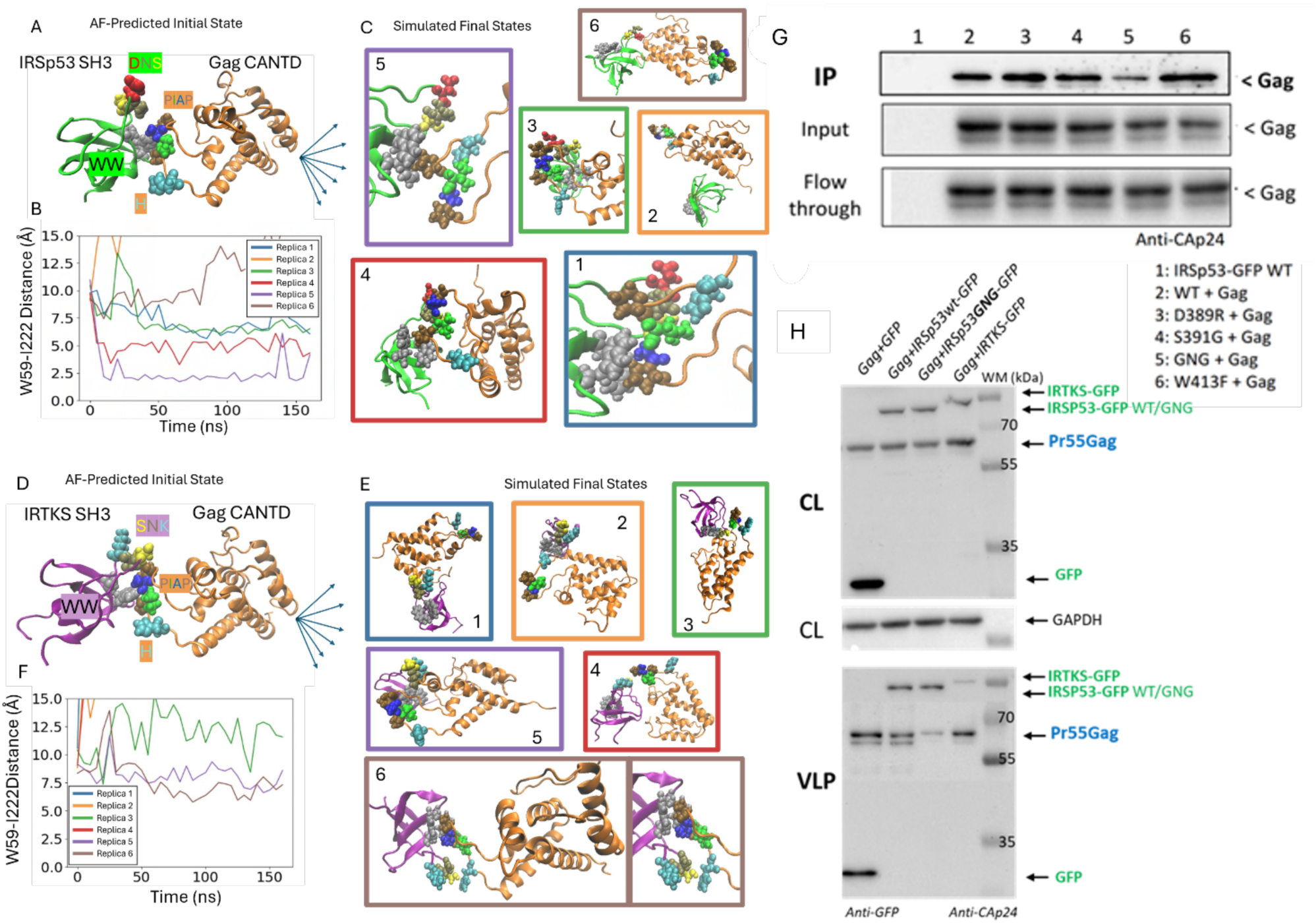
Interactions between the IRSp53 SH3 and the Gag CANTD domains. (A) The initial state for molecular dynamics was the Gag CANTD from previous work(4) and PDB 2M8L for the SH3 domain fit to the AlphaFold-predicted structure of the interface. (B) We ran 6 replicates and tracked the W59-1222 distance to measure stability. The colors correspond to the colors outlining the final snapshots in (C). There are two main modes of interactions corresponding to interactions with different prolines in the PIAP and some dissociation. The DNS domain is heavily involved in the stable interfaces. (D) The initial state for molecular dynamics was the gag CANTD from previous work(4) and PDB 2KXC for the SH3 domain fit to the AlphaFold-predicted structure of the interface. (E) We ran 6 replicates and tracked the W59-1222 distance to measure stability. The colors correspond to the colors outlining the final snapshots in (F). There are two main modes of interactions corresponding to interactions with different prolines in the PIAP and some dissociation. The SNK domain is heavily involved in the stable interfaces. (G) Immunoprecipitation results revealed by western blots (anti-CAp24) confirm that DNS mutation in the IRSp53 SH3 domain results in a much weaker interaction between Gag and IRSp53 than the WT. IP is Gag immunoprecipitated with IRSp53-GFP thanks to the GFP-Trap kit; Input is the total Gag proteins; the Flow-through is the Gag proteins that are not bound to IRSp53-GFP-beads complex during the experiment. These are representative western blots of 3 independent IP experiments. (H) Western blots showing Gag-VLP release from transfected HeLa cells expressing Gag and either exogenous GFP, IRTKS-GFP, IRSp53-GFP WT or GNG mutants, showing that Gag-VLP release is impaired by the IRSp53-GFP GNG mutant. WM is the weight marker in kDa.

We then investigated the role of other specific amino acids in the SH3 domain. In Figure 3D, we show some that took away the D and S amino acids of the DNS domain and replaced them with a glycine (GNG mutant). This could provide not only further support that the SH3 domain is the interaction site but also the molecular mechanism discovered with the MD. We sought to confirm our prediction that IRSp53 and Gag is due to a direct interaction between the SH3 domain of IRSp53 and the CANTD of Gag using direct mutagenesis of the DNS IRSp53 motif. Then we performed immunoprecipitation experiments with IRSp53 mutated forms. Our experimental results in cells show a loss of interaction between the GNG IRSp53 mutant and Gag (Figure 3G) as compared to single mutations in D389R or S391G. Following this, we analyzed the effect of overexpression IRSp53 or IRTKS or IRSp53 GNG mutant in Gag-VLP release (Figure 1E). Our experimental results reveal that the overexpression of IRSp53 GNG mutant acts like a dominant-negative by reducing Gag-VLP release as compared to WT IRSp53-GFP, while IRTKS-GFP has no effect on Gag-LP release.

### IRSp53 SH3 Domain Alters Gag MACANTD Conformation as in RNA-Bound Gag

Experimentally, to analyze which domain of HIV-1 Gag protein was involved in the interaction with IRSp53, we tested different known Gag mutants (Figure 4A). Gag was tagged by mEos2 to visualize Gag expression in transfected cells. Gag-dHBR (MA-highly basic region) was reported to abrogate Gag plasma membrane targeting through the phospholipid PI(4,5)P2 (Ono et al, PNAS 2004(*46*)), Gag-WM was reported to impair Gag multimerization through CA-CA interaction(*24, 47*)), Gagdp6 impaired Gag-VLP budding(*48*) and MACA was truncated for the NC-sp2-p6 domains of Gag and loose the capacity of creating assembly platform at the cell surface (Floderer et al, 2018(*49*)). Armed with these Gag mutants, we performed an immunoprecipitation of endogenous IRSp53 and revealed by Western Blot (anti-Cap24) if any WT Gag or mutants were pulled-down. The experiment shows that the MA and CA domain of Gag were relevant interaction domains (Figure 4B). Moreover, the ratio of IP over whole cell lysate, shows that most probably, Gag needs to assemble at the cell membrane and multimerize to interact with IRSp53, as no IP was detected with the double mutant dHBR/WM.

**Figure 4:**
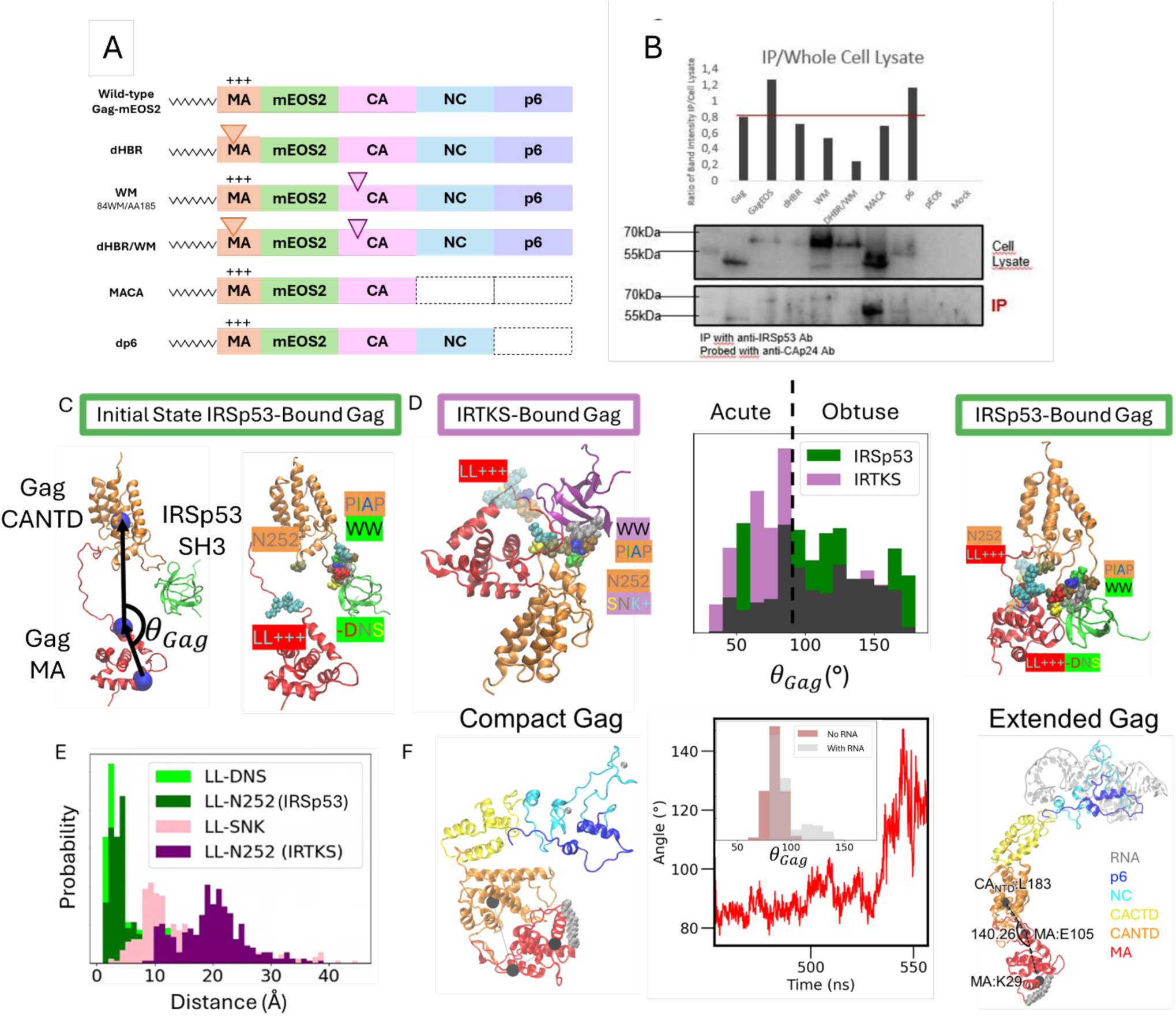
The effect of SH3 domains and RNA on Gag MACANTD conformations. (A) HIV-1 Gag mutants defectuous in assembly and budding used in the study. WT Gag(i)mEOS2, dHBR, having a deletion in the highly basic region in MA; WM holding a mutation in CA reducing Gag multimerization; MACA is a Gag depleted of its NC-p 6 C-ter domains preventing assembly; dp6 Gag is depleted of its p6 domain that recruits ESCRT for particle budding. These Gag mutants were described previously in Floderer et al., 2018). (B) Immunoprecipitations and Western blots showing the cell lysates containing Gag mutants and the results of the interaction with the endogenous IRSp53 (IP) thanks to an anti-CAp24 revealing Gag. The graph shows the ratio between the IRSp53 found in the IP and the cell lysates for Gag content. Simulations of the Gag MACANTD bound with SH3 domains start from the configuration shown in (C) with the important residues and motifs shown for IRSp53 The angle *θ_Ga_*_g_ quantifies how extended the Gag MACANTD is. The lower linker domain is LL and notation for other motifs is described in the text. (D) IRTKS-Bound Gag lacks contacts with the lower linker domain found in IRSp53-Bound Gag that lead to extended conformations as quantified by the distribution of *θ_Ga_*_g_. (E) Quantification of the contacts with the lower linker shown in (D) by the closest distance between the lower linker and relevant contact sites. (F) The full Gag polyprotein undergoes a transition from Compact to Extended upon binding RNA. The angle *θ_Ga_*_g_ shows a change like the change brought on by IRSp53 binding. The time series shows one example of this spontaneous transition and the distribution shows it never occurs without RNA, but the distribution for the with RNA case is likely undersampling the extended state.

To fully understand the role of the MA domain in this interaction, we also performed simulations using a Gag MACANTD construct as shown in Figure S3, where the Gag was locked into a “multimerized” conformation, which is an extended conformation. We found that both IRTKS and IRSp53 SH3 domains fit nicely between the MA and CANTD and can interact with both while sitting in the pocket between them. Between these results and our results with just the CANTD of Gag we had no evidence that there should be any difference between IRSp53 and IRTKS. Thus, we decided to simulate the Gag MACANTD construct with no constraints, which we hypothesized would be more like the Gag before it assembles into the lattice.

We ran 4 MD replicas for at least 450ns for IRTKS and IRSp53 (total of 3.5 microseconds) and observed that the Gag MACANTD sampled different conformational states based on whether it was bound to IRTKS or IRSp53. We quantified this using the angle, *θ_Ga_*_#_, which is shown in Figure 3. This angle quantifies how extended is the conformation of the MACANTD Gag, which we hypothesize is important for Gag to make both MA-MA contacts and CATND-CANTD contacts necessary for lattice formation. IRSp53-bound Gag was in more extended conformations than the IRTKS-bound Gag as shown in Figure 4D. For the first time, this gave us a possible explanation for the experimentally observed differences between the two I-BAR proteins(*15*). We also quantified the amino acid residues which appeared to be responsible for this. We noted that the DNS domain of IRSp53 was interacting electrostatically with the 3 Lysines found on the lower linker domain by MA (Lys111, Lys112, Lys113). It seems that this connection along with the WW-PIAP interaction described earlier effectively superseded the linker as the connection between MA and CANTD leading to more extended, less flexible conformations (see Figure 4E). The MA and CANTD also tended to interact directly through a hydrogen bond formed between the lower linker and ASN252 in the CANTD (see Figure 4E). In contrast, IRTKS does not have a DNS domain. Instead, it has an SNK domain (Ser22, Asn23, Lys24) meaning that the electrostatic attraction with the lysines on the lower linker of Gag become an electrostatic repulsion and that all the conformation stabilizing interactions in the IRSp53-bound Gag do not exist in the IRTKS-bound Gag (see Figure 4D and E).

We desired to obtain further evidence that these extended conformations were relevant to the actual behavior of Gag. To do this we further analyzed previous simulations of Gag bound to RNA(*9*). It is known that RNA activates Gag by changing the conformation and so we wanted to see if these conformational changes to the MACANTD construct could also be triggered by RNA. This is shown in Figure 4F as binding RNA triggers a spontaneous increase in the angle which is very similar to the difference between IRTKS and IRSp53-bound Gag conformations. Without RNA we see almost entirely acute angles, matching more closely with the IRTKS case.

### Rationally Designed Gag Mutants are not Conformationally Extended by IRSp53

The results described above suggest that the difference between IRTKS and IRSp53 is not in the ability to bind Gag but the ability to activate it for assembly. This explains why knocking out IRTKS did not show the same effect and why IRTKS, which also binds Gag (experimentally and *in silico* as we demonstrated), cannot compensate for IRSp53 being knocked out. However, we still sought more evidence for the molecular mechanism we have proposed. Thus, we ran the MD simulations of IRSp53 bound Gag three more times, with mutations in Gag that we hypothesized could change the observed Gag conformations. In the first mutant, we removed the polyproline helix which binds the TRP prong of the SH3 domain by replacing the two prolines in the PIAP motif with alanine. This is referred to as AIAA Gag (Figure 5A-D). In the second mutation, we removed the interaction between the DNS motif and the lower linker lysines, which seemed to be allowing IRSp53 to fix the Gag conformation. We mutated all 3 lysines to alanine and thus we refer to this mutant as 3K3A Gag (Figure 5E-H). Finally, we mutated the 3 glutamic acids on the MA domain we found were interacting in the multimerized state in Figure S3 and we refer to this mutant as 3E3A (Figure 5I-L).

**Figure 5:**
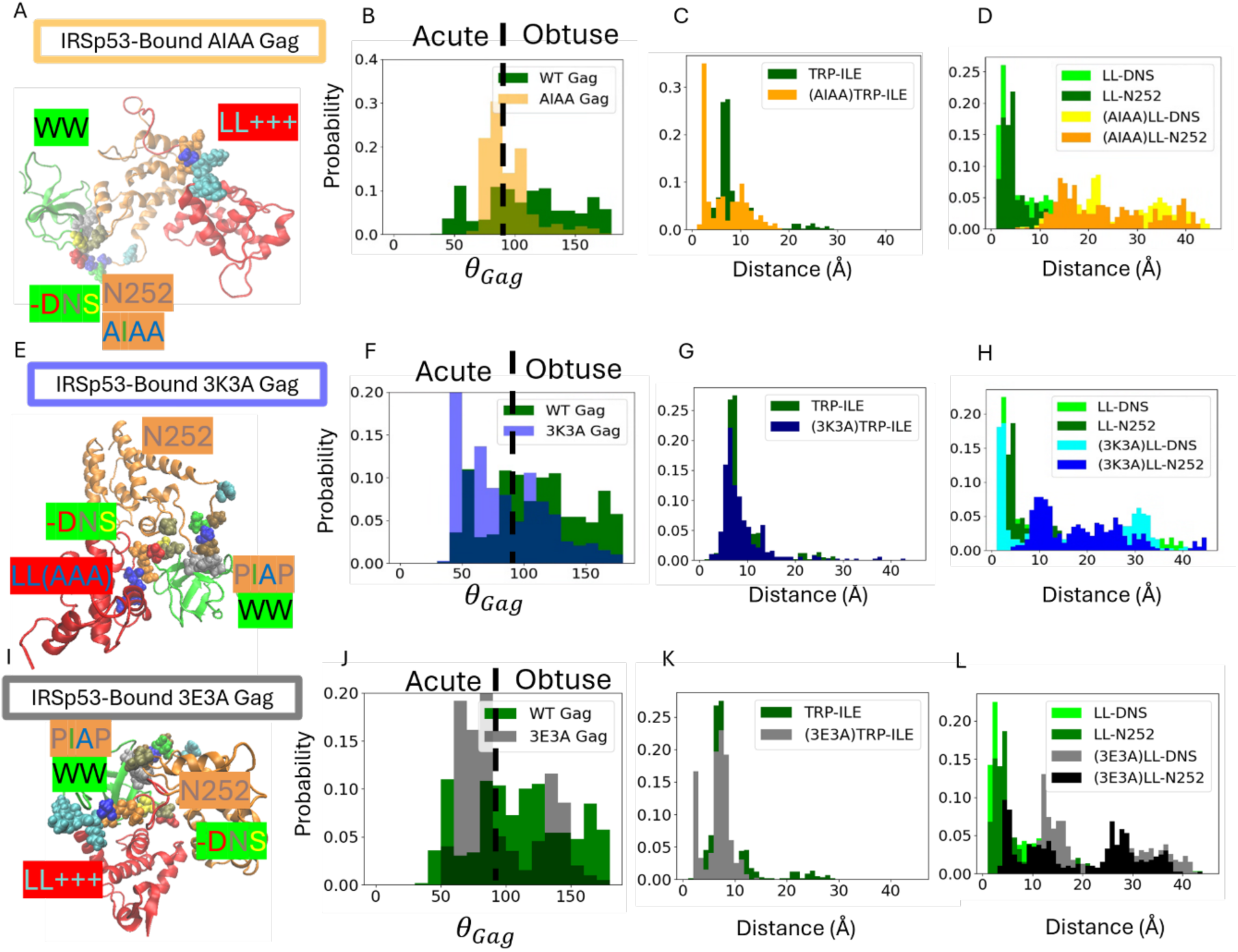
Mutated Gag does not extend in the presence of IRSp53. (A) A snapshot shows AIAA Gag (P221A, P224A) does not make the correct contacts with IRSp53 to create extended conformations like WT Gag. (B)The distribution of angles shows much less extended conformations. (C) The distance between the I of AIAA and the TRP prong (W59 of IRSp53 SH3) are altered due to removing the polyproline domain of Gag. This leads to altered interactions with the lower linker (LL) domain of Gag in (D), which lead to less extended conformations. (E) A snapshot shows 3K3A Gag (K111A, K112A, K113A) does not extend in the presence of bound IRSp53 due to a lack of interaction between N252 on the CANTD and the lower linker. This is quantified through the angle distribution in (F). The TRP prong interaction with the PIAP of Gag is still intact as shown by the W59-I222 distance in (G), but the N252-LL distance in (H) is altered due to the inability of N252 to hydrogen bond with the residues 111-113 on the lower linker once they were mutated from K to A. Lower linker contacts with the DNS motif of IRSp53 SH3 exist to some extent. (I) A snapshot shows 3E3A Gag (E104A, E105A, E106A) does not extend in the presence of bound IRSp53 due to a lack of interaction between N252 on the CANTD and the lower linker. This is quantified through the angle distribution in (J). The TRP prong interaction with the PIAP of Gag is still intact as shown by the W59-I222 distance in (K), but the LL-N252 and LL-DNS distances in (L) are much higher.

We analyzed these simulations using the same criteria as we did previously for WT Gag with IRSp53 and IRTKS. We found that for all three of our mutants the distribution of Gag angles (Figure 5B, F, and J) showed more acute than obtuse angles, meaning that our hypothesis that these Gag mutants would not be moved towards active conformations in the presence of IRSp53 was accurate, at least *in silico*. We observed that the AIAA Gag did not interact in the same way with the TRP prong (Figure 5C) leading the DNS, lower linker, and N252 to all be misaligned and unable to stabilize extended conformations (Figure 5D). For the 3K3A Gag, the TRP prong interacted normally with the polyproline helix as expected (Figure 5G). We also observed an interaction between DNS and the lower linker that was more like the wild type than we expected given the mutation of all three lysines to alanine (see the cyan bimodal distribution in Figure 5F). It is worth noting that in the case of IRTKS this interaction is fully repulsive between the Lys24 of the SNK and the three Lysines of the lower linker, while for 3K3A Gag the electrostatic interaction is neutral. The main difference for the 3K3A Gag seemed to come in the inability of N252 to form hydrogen bonds with the lower linker after the mutation of the lysines to alanines (Figure 5F). For 3E3A Gag, we did not directly mutate amino acids involved in the interaction and thus we thought there might not be a large difference in Gag conformations. However, Figure 5J shows the conformation is less extended than the WT and that the lower linker interactions are disrupted (Figure 5L) while the TRP prong interactions are not (Figure 5K). It seems that mutation of the three glutamic acids changes the local electrostatic environment enough to alter the interactions of the 3 Lysines and thus Gag conformation.

### Rationally Designed Gag Mutants Display Assembly and Budding Defects in vivo

As a final step, we sought to confirm in cells that these Gag mutants, which we showed *in silico* should not be activated by IRSp53, would have the same or similar phenotype to what was observed when IRSp53 was knocked out completely. This would serve to help validate our hypothesis that altering Gag conformation is a key role of IRSp53 and one that cannot be rescued by IRTKS or other I-BAR proteins.

We expressed fluorescent chimera (Gag-i-GFP) of WT Gag and the different mutants (AIAA, 3K3A, 3E3A) in HeLa cells. We first imaged cells transfected by WT Gag and the different mutants (Figure6A) using TIRF microscopy to increase signal of the cell surface attached to the coverslip, allowing for identification of (individual) on-going assemblies. Whereas WT Gag transfected cells exhibit mostly sparse, individual on-going assemblies at their surface, the three mutants show patches/clusters of either on-going assemblies or aggregated Gag proteins. Using image segmentation, we could determine the cell surface proportion occupied by these on-going assemblies/Gag aggregates (N≥21 different cells for each condition). Figure 6B shows increasing surface coverage of the three mutants (AIAA<3E3A<3K3A) versus WT Gag, suggesting either more efficient assembly, accumulation of non-fully assembled particle at the cell surface, or less efficient particle release from the cell surface. We then measured the average fluorescence intensity per surface of each segmented individual and clusters of on-going assemblies and normalized this fluorescence to the whole cell surface averaged fluorescence in order to have an idea of the distribution of local enrichment in Gag proteins in the identified clusters. Figure 6C shows these distributions of each segmented area for each mutant (5100(3E3A) < n < 6900(WT) segmented clusters and N≥21 different cells for each condition). AIAA exhibits the highest median value and, more interestingly, the widest distribution of Gag enrichment in segmented clusters. On the contrary, WT Gag shows the narrowest distribution for Gag enrichment in segmented clusters. This narrow distribution suggests that most of the identified clusters in cells represent one single on-going/finished assembly site, which Gag enrichment compared to the cell peaks at 1.2x. Interestingly, the 3K3A mutants also peaks at 1.2x enrichment with a (slightly larger but) narrow distribution, suggesting a large presence of individual on-going assembly. However, the surface coverage of 3K3A mutant is twice the one of the WT Gag. Finally, the 3E3A mutant generates clusters which characteristics are in between WT Gag (slightly higher enrichment peak,1.3x) and AIAA for the discrepancy of the distribution. The main advantage of categorizing the mutants by this approach is the high statistics (n>5100 assembly sites/Gag clusters, in N>21 different cells).

**Figure 6:**
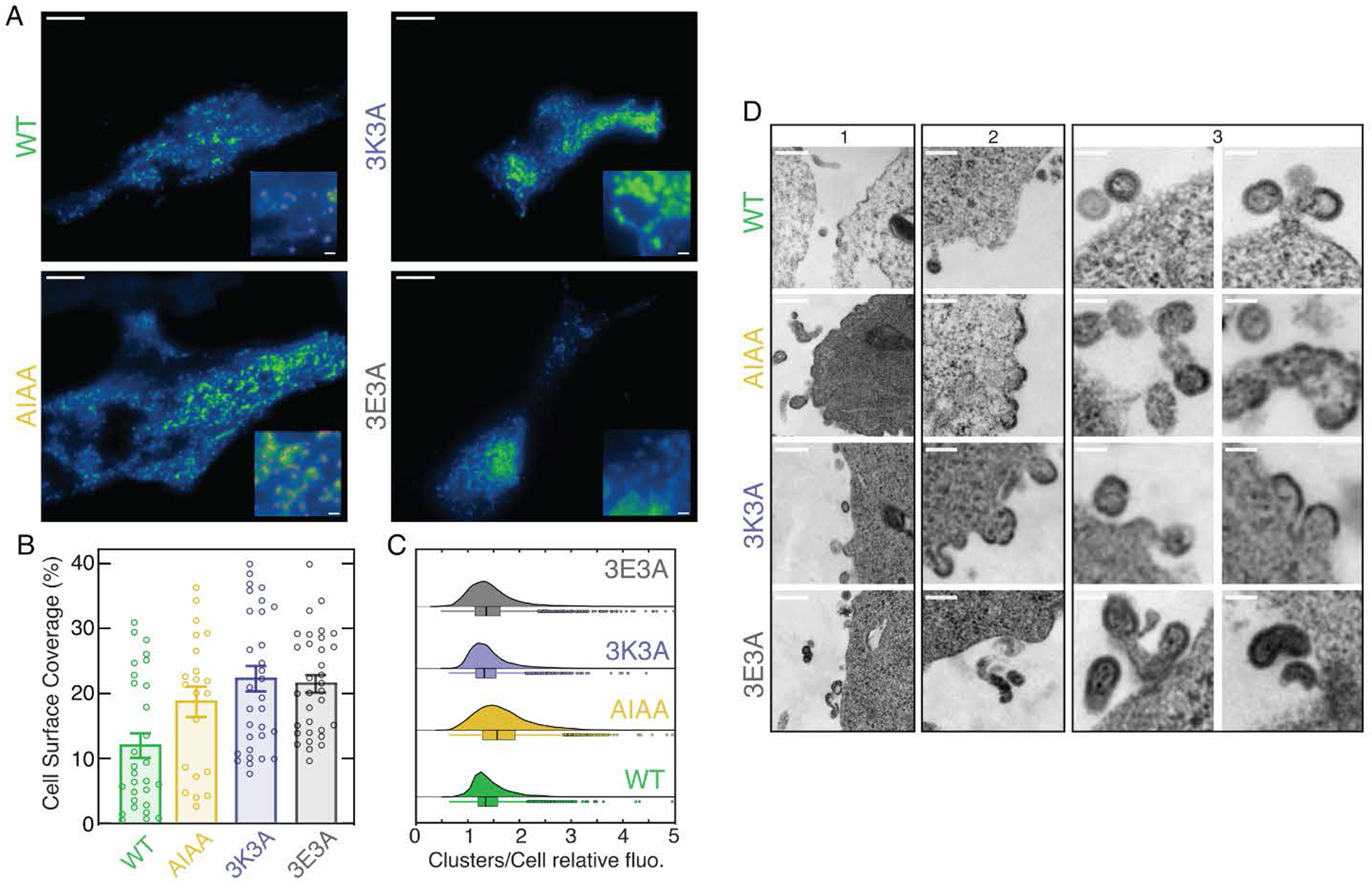
Budding of computationally designed Gag mutants is stalled at the plasma membrane. A-TIRF microscopy images of HEK293T cells expressing WT Gag and the different computationally designed MACA mutants AIAA, 3K3E, and 3E3A (scale bars are 5µm and 500 nm in the zoomed insets). B-Cell surface coverage (in %) of the sum of identified clusters after segmentation (for segmentation examples, see purple surrounded areas in WT and AIAA insets TIRF images). C-Distribution of the relative mean fluorescence intensity of segmented clusters vs mean fluorescence intensity of the rest (non-cluster areas) of the cell. D – Electron micrographs of WT Gag and the different computationally designed mutants. Scale bars are 500 nm (panels 1), 200 nm (panels 2) and 100 nm (panels 3).

In addition, electron microscopy, albeit lower statistics, provides better insight into the structures of each ongoing assembly site. Figure 6D shows typical snapshots at increasing magnification (from panel 1 to panel 3) for the different Gag. At the highest magnification (panel 3), WT Gag electron micrograph shows fully assembled spherical virus-like particles with a diameter of ∼130nm. The 3K3A mutant also exhibits spherical VLPs (see panel 1 and 3) and numerous on-going assemblies with a partial coverage in Gag proteins (panel 3). By contrast, AIAA, as well as the 3K3E mutant shows a large coverage of the membrane with Gag proteins (panel 2), accounting for the largest surface coverage observed in the TIRF microscopy, and grapes/clusters of particles more or less spherical (panel 3), budding from the cell surface.

Overall, the results presented above support the hypothesis that the Gag mutants exhibit assembly and budding defects that appear to result from impaired Gag–IRSp53 interaction.

## Discussion and Conclusions

We initiated the intellectual thread of this article starting from no information on whether IRSp53 was directly interacting with Gag and what the molecular origin of the interaction might be. AlphaFold predictions suggested that the SH3 domain of IRSp53 was interacting with the CANTD of Gag. We were able to interpret these predictions using our understanding of polyproline domains and specific relevant structures that were available in the HIV-1 literature and PDB more widely. These structures were most likely what AlphaFold originally trained on and in this way the predictions could be considered a type of interpretable machine learning. This understanding even allowed us to predict how the AlphaFold prediction may be deficient. Using comprehensive MD simulations, we were able to determine how the interface rearranged from the AlphaFold structure and the DNS/SNK domains of IRSp53 and IRTKS become a stabilizing force. Our subsequent experiments confirmed the importance of the non-I-BAR parts of the IRSp53, especially specific residues in the SH3 domain, some of which were only implicated by the MD and not by AlphaFold. Additionally, they showed that the MACA domain of Gag and Gag multimerization on membranes are relevant to the Gag-IRSp53 interaction. This motivated us to extend the MD simulations to include the MA domain of Gag. We saw that when Gag was held in the lattice-like conformation, the SH3 domains of both IRSp53 and IRTKS fit well between the MA and CANTD, interacting with residues on both domains. When we released Gag from the multimerized conformations, we saw that IRSp53 SH3 and IRTKS SH3 had different impacts on Gag conformation. IRSp53 had a much greater tendency to bias Gag into extended conformations which also look like the conformations induced by RNA binding. Since it is known that these RNA-bound conformations are more active for Gag lattice assembly, we hypothesized that Gag which binds IRSp53 before multimerization would become more active for assembly, while Gag bound to IRTKS would not. This behavior could catalyze Gag assembly and the integration of IRSp53 into the periphery of Gag clusters, but not IRTKS. In turn, this could provide an explanation for why IRTKS cannot rescue Gag budding when IRSp53 is knocked out.

We then used the previous MD simulations to design two Gag mutants that would not change conformation in the presence of IRSp53 and one that would affect the binding of SH3 domains in the multimerized state. We next verified the effect of these mutants with further MD simulations. Experimentally, all of these mutants led to a substantial accumulation of Gag at the plasma membrane, indicating that Gag assembly at the plasma membrane was impaired in its ability to progress to the budding of immature viral particles. TEM images further revealed that even the viral particles that would bud were malformed. These results confirmed that the Gag mutants were assembling but had a significant defect in budding. We assert this provides evidence that Gag assembles with IRSp53 and that this is helps to provide the membrane curvature necessary for virion budding. This assembly requires IRSp53 to interact directly with Gag through its SH3 binding to the MACANTD domain of Gag. It appears that this interaction occurs before Gag monomers join the lattice since the SH3 domain of IRSp53 activates the conformational state. IRTKS SH3 and Gag mutants do not have this conformational activation and therefore exhibit the defective downstream budding. It may also be that Gag bound to IRSp53 is more readily integrated into the lattice, allowing IRSp53 to more readily bind to the lattice periphery. Such an effect would likely require coarse-grained molecular dynamics to better understand and these studies are underway.

## Acknowledgements

Research reported in this publication was supported in part by the National Institutes of Health (NIH) through the Behavior of HIV in Viral Environments (B-HIVE) Center of the National Institute of Allergy and Infectious Diseases (NIAID) under award number U54AI170855 (50%) and in part by the NIH NIAID award number R01AI178850 (50%) (both to G.A.V.). C.W. acknowledges the support of an NIH Ruth L. Kirschstein National Research Service Award Postdoctoral Fellowship under award number F32AI186429. The content is solely the responsibility of the authors and does not necessarily represent the official views of the NIH. Computational resources were provided by the Research Computing Center and the NIH-funded Beagle-3 computer (NIH award 1S10OD028655-01). This work was supported in part by high-performance computer time and resources from the DoW High Performance Computing Modernization Program. We thank Giorgio Scita and his laboratory (FIMO, Milan, Italy) for giving us access to the IRSp53-GFP mutants (234, IMD and dIMD). We also thank the Abbelight company and MRI core facility in Montpellier for the use of the TIRF-Microscope, and further thank the University of Tours for electronic microscopy core facility.

## Contributions

CW and JB for AlphaFold predictions and interpretation. CW and PB for Molecular Dynamics modeling. PM and KB for directed mutagenesis on IRSp53-GFP and Gag/Gag(i)GFP plasmids, molecular and cellular biology. PM, KB and CP for IP and Western Blots. RG for TIRF-Microscopy imaging. JBG for TEM imaging. CW and DM for first draft writing. CW, GV, CF and DM for manuscript editing and grant raising.

## Materials and Methods

### Computational

#### AlphaFold

We used the AlphaFold webserver which runs AlphaFold3(*51*) to predict the structure of Gag and IRSp53. The sequences can be found in the SI. All structures were downloaded and visualized using VMD(*52*), with the first and most confident of the 5 structure predictions being deemed as a reasonable candidate based on similarity with published structures.

#### Structure Preparation and Mutation

For the simulations with just the SH3 domain we started by aligning the known structure of Gag CA (PDB 2M8L(*53*)) with the alphafold predicted structure using only the PIAP domain residues. We then did the same using the structure of the IRSp53 SH3 domain using the DGW 44-46 residues that end with the TRP residue predicted to be inserted into the PIAP by alphafold. The CA and IRSp53 proteins were cut into only the CANTD and SH3 domains (see SI Figure S4 for details on sequences used). The process was repeated for IRTKS using the structure from PDB 2KXC using the same DGW domain.

For all simulations with the Gag MACANTD we started from the structure of a Gag 4-mer from our previous work(*4*). We then took the structure from one of our MD replicas with and fit that onto the Gag structure using the PIAP (221-224) motif residues only. We then combined the structure from the 4-mer with the now-aligned SH3 domain to create a stable interface. The process was repeated with IRTKS.

Alignments were performed using MSBack utilities(*54, 55*) and sidechain clashes were fixed with Flowback(*56*).

Mutations were performed by taking the same initial structures, changing the sequence and then solving only the new sidechains and any potential clashes with flowback.

#### MD Simulations with SH3 Domains

We ran several sets of MD simulations and the initial states of the proteins were prepared as described above. We then solvated and equilibrated the systems as described here. TIP3P water was added to the system to generate the solvated structure in a box whose size varied based on the specific system(*57*). More details can be found in the SI. We neutralized the system with a NaCl concentration of 150 mM. The periodic boundary condition was active across the three dimensions of the simulation box. The CHARMM36 force field was used with the particle mesh Ewald sum method for long-range electrostatic interactions(*58*). After energy minimization with the steepest descent algorithm, the system was pre-equilibrated at 1 atm and 310 K (the constant *NVT* ensemble followed by the constant *NPT* ensemble). All production runs were performed with a 2 fs time step in the constant *NPT* ensemble using Parinello–Rahman pressure coupling and Nose-Hoover temperature coupling.

The first set of simulations involved only the SH3 domains and the Gag MACANTD. Each of the 6 production runs per SH3 domain were run for 160 ns. Thus, the total time was slightly less than 2 microseconds.

We ran 4 sets of simulations included unrestrained Gag MACANTD. Two of the sets involved the IRSp53 and IRTKS SH3 domains. The other two had IRSp53 with a mutated form of Gag. All these simulations were run as 4 replicas for at least 400 ns each totaling 1.6 microseconds per set and nearly 7 microseconds total.

The last set of simulations involved both the SH3 domains and the Gag MACANTD, which was restrained to simulate the multimerized state in the Gag lattice (see SI for full detail). We ran 4 replicas with the IRSp53 SH3 domain and 4 replicas with IRTKS SH3 domain for nearly 1 microsecond each totaling over totaling over 3.5 microseconds for each SH3 domain. These simulations were much faster than the unrestrained version since the restraints allowed us to use a smaller box without worrying about the PBC since the Gag conformation did not change throughout.

### MD Simulations with RNA

#### Analysis

Throughout the paper we measure the conformation of the Gag using the angle, *θ_Ga_*_g_. We define this angle using the positions of Lys29, Glu105, Leu183. Specifically, the arccos of the dot product between the unit vector starting at the alpha carbon of Lys29 and pointing to the alpha carbon of Glu105 and the unit vector starting at the alpha carbon of Glu105 and pointing to Leu183.

Other distances between residues reported in the paper reflect the closest distances between any pair of two atoms in the residue or set of residues. The lower linker domains were defined to include residues 111-116 which includes the three lysines as well as Ala114 and Gln115,116.

### Experimental

#### Material and Methods

(BIOLOGY)

#### Antibodies

A rabbit polyclonal anti-IRSp53 antibody (Merck Millipore), a mouse monoclonal anti-CA (NIH AIDS Reagent Program), a rabbit polyclonal anti-GFP (Invitrogen) were used in this study.

**Table 1:** Antibodies used in this study.

|  |  |  |
| --- | --- | --- |
| Anti CAp24 mouse | NIH clone 183H125C | 3537 |
| Anti eGFP rabbit | Life technologies | A11122 |
| Anti GAPDH rabbit | Sigma | G9295 |
| Anti Rabbit-HRP | Agilent | P0217 |
| Anti Mouse -HRP | Agilent | P0260 |

#### Plasmids

The plasmid expressing HIV-1 codon-optimized Gag (pGag(myc), named pGag) and the pGag(i)GFP located between MA and CA were used in this study. Mutants were made in both plasmid expressing Gag and Gag(i)GFP using the mutagenesis kit QuikChange II XL Site-Directed Mutagenesis Kit (Agilent200521). The WT IRSp53-GFP expressing plasmid (Inamdar et al., 2021(*15*)) was mutated in the GNS motif of the IRSp53 SH3 domain using the QuikChange II XL Site-Directed Mutagenesis Kit (Agilent200521). All plasmids were amplified at 37°C in E.Coli and purified using the Xtra maxi plus EF kit (Macherey-Nagel 740426) and sequenced using MWG Eurofins facilities to confirmed mutations in pGag(myc), pGag(i)GFP and pIRSp53-GFP.

For pGag and pGag(i)GFP, substitution mutations were introduced in MA to replace a triplet of Glutamic acid (E104-E106) by a triplet of Arginine (A104-A106). Another substitution mutations were introduced in MA to replace a triplet of lysine (K111-E113) by a triplet of Arginine (A111-A113). For pGag, substitution mutations were introduced in CA to replace P90AIP93 by A90AIA93 (named AAIA).

For IRSp53-GFP, 4 mutants were generated by substitution: Aspartic acid 389 was replaced by Arginine (D389R), Serin 391 was replaced by glycin (S391G), Tryptophan was replaced by Phenylalanin (W413F) and a double mutant by replacing Aspartic acid 389 and Serin 391 by a Glycin. (GNG).

#### Cell culture and transfection

Human embryonic kidney cells (HEK 293T-ATCC-CRL-1575TM) were maintained in Dulbecco’s Modi fied Eagle’s Medium (DMEM, GIBCO) and human HeLa cells (ATCC-CRL-CCL-2). All cell lines were tested negative for mycoplasma thanks to a MycoAlert Mycoplasma Detection Kit (Lonza Bioscience) done every month. Media was supplemented with 10% fetal bovine serum (FBS, Dominique Dutscher) and complemented with sodium pyruvate and antibiotics (penicillin-streptomycin). Cells were grown at 37°C in a 5% CO2 atmosphere. Transfection was performed by using the calcium phosphate precipitate method on HEK293T cells and the JetOptimus transfection for HeLa cells using the manufacturer’s protocol. The number of cells and amount of transfected plasmid varies in function of the experiments (IP, Western blot or Microscopy). The cell medium was replaced by fresh medium 6 hr post-transfection and the experiments were performed 24 to 48 hr post-transfection.

#### Immunoprecipitation assay

Based on different plasmids conditions, HEK293T cells (2.8.10e6 cells) were transfected with pIRSp53-GFP plasmid alone or with pGag WT or pGag mutants, or with eGFP plasmid alone or with pGag WT or mutants (10µg of total DNA). The cell medium was replaced 6 hr post-transfection. After 24h post transfection, cells were detached using EDTA 1mM and the pellet was lysed 30min on ice in lysis buffer prepared using the manufacturer’s protocol supplemented with Complete™, Mini, EDTA-free Protease Inhibitor Cocktail (Sigma 11836170001). Lysates were pelleted 10min at 4°C at 17000g and diluted in supplemented lysis buffer. 550µL of lysates were added to 25µL magnetic agarose bead covalently bound with an anti-Green Fluorescent Protein (GFP) Nanobody (VHH) (ChromoTek GFP-Trap® Magnetic Agarose Kit (gtmak-20). The complex was incubated for 1h30 on the tube rotator at 4°C. The samples were then washed four times with the wash buffer, followed by addition of 80µL of 2X Laemmli’s buffer to the beads. The samples were denatured at 95°C for 5 min and then processed for Western blot.

#### Western blot and analysis

About 50 mg of each protein (intracellular in cell lysates) samples or 20 mL of purified VLP samples were mixed with SDS loading dye, deposited, and resolved on a 10% SDS-PAGE gel. The gels were then transferred on to polyvinylidene PVDF membranes (Amersham). Immunoblotting was per formed by incubating the membranes overnight with primary antibody at 4°C, and 2 hr with horse radish-peroxidase (HRP)-conjugated secondary antibody at room temperature. The Western blot signals were detected using ECL Prime/ECL Select substrate (Amersham) and images were taken using ChemiDoc (Bio-Rad).

#### VLP purification and quantification

After 24 or 48 hr post-transfection, cell culture supernatants containing Gag-VLPs were collected, filtered through a 0.45 mm filter, and clarified at 800 g for 5 min at 4°C. The supernatant was then purified by loading it on a cushion of 25% sucrose in TNE buffer (25 mM Tris-HCl, 4 mM EDTA, and 150 mM NaCl) and ultracentrifuged at 100,000 g for 1 hr 30 min at 4°C in an SW41Ti rotor (Beckman Coulter). Dry pellets were resuspended in TNE buffer at 4°C overnight. Gag-VLP release was estimated by performing anti-CAp24 immunoblot and by quantifying Gag signal in the blots using Fiji software as described in Floderer et al., 2018(*49*). The calculation for Gag-VLP release is: % of Gag in VLP = Gag released / (Gag released + Gag intracellular normalized to GAPDH)

#### TIRF-Microscopy

HeLa or HEK293T cells expressing HIV-1 Gag/Gag(i)GFP or mutants were cultured on poly-l-lysine (Sigma) coated 25 mm round #1.5 coverslips (VWR) for 24h prior observation then fixed using 4% PFA in PBS for 15 min at room temperature, rince 3 times with PBS and imaged in PBS using a glass bottom culture Fluorodish using a Total Internal Reflexion Fluorescence-Microscope (Abbelight). Images were taken with a sCMOS camera, using a NA = 1.45, 100 objective and a 488 laser.

#### Transmission electron microscopy

Transfected HEK293T cells were fixed in 4% paraformaldehyde and 1% glutaraldehyde in 0.1 M phosphate buffer (pH 7.2) for 48 hr, washed with PBS, post-fixed in 1% osmium tetroxide for 1 hr, and dehydrated in a graded series of ethanol solutions. Cell pellets were embedded in EPON resin (Sigma) that was allowed to polymerize at 60°C for 48 hr. Ultrathin sections were cut, stained with 5% uranyl acetate and 5% lead citrate, and deposited on collodion-coated EM grids for examination using a JEOL 1230 transmission electron microscope.

**Table 2:**
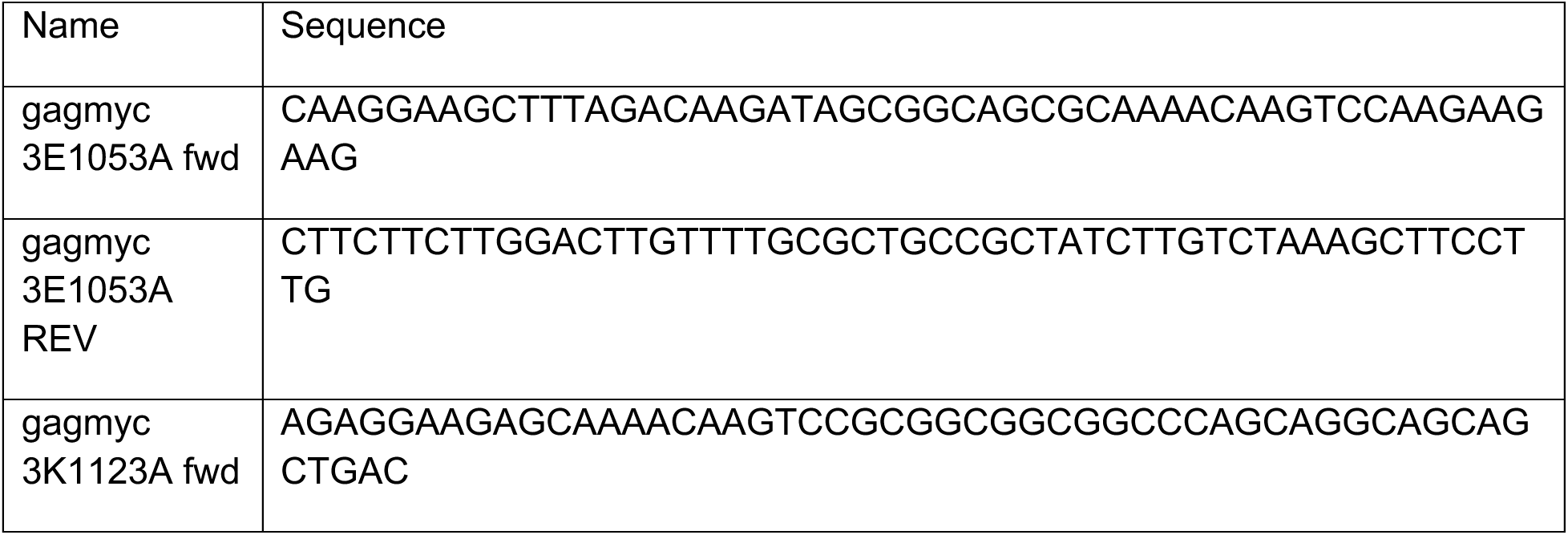
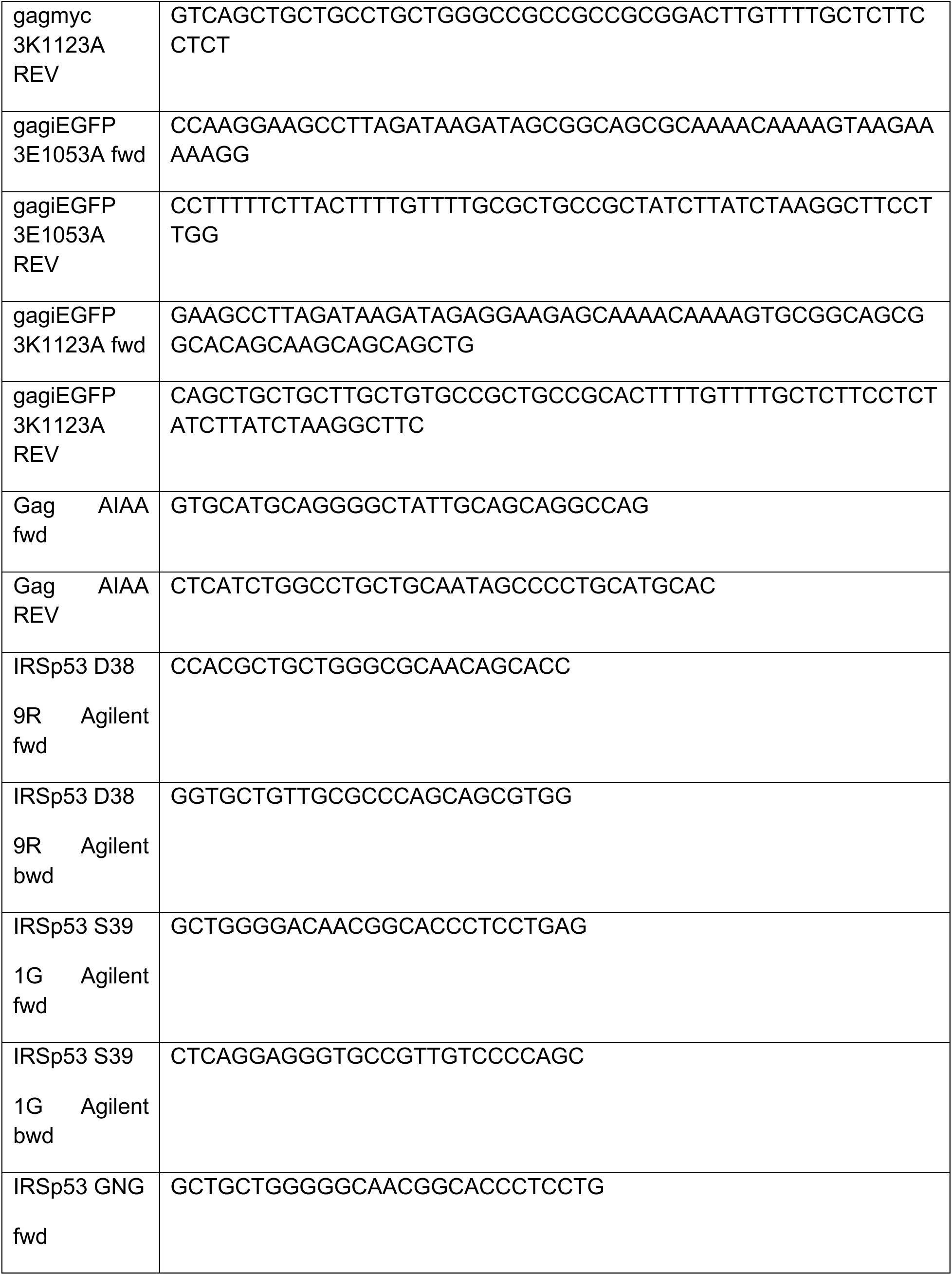
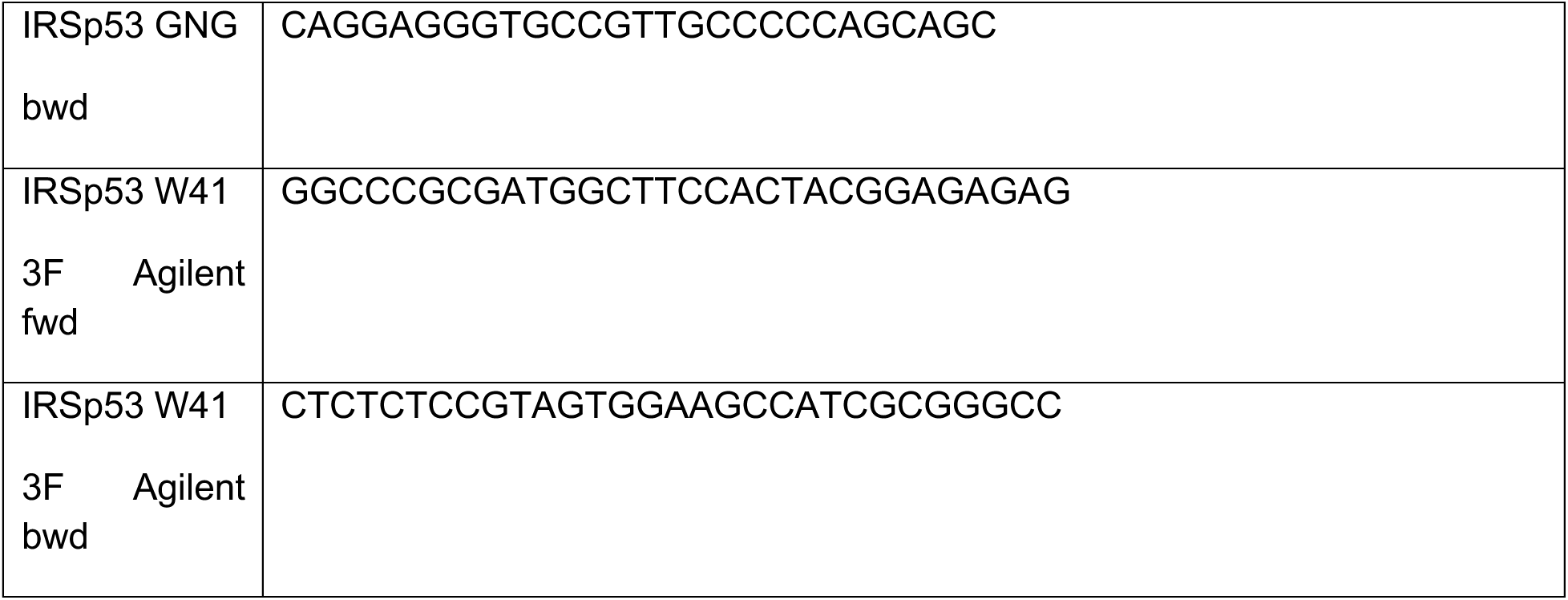
Primers used to generate Gag and IRSp53 mutants.

## References

1. E. O. Freed, HIV-1 assembly, release and maturation. Nat Rev Microbiol 13, 484–496 (2015).

2. A. Tan, A. J. Pak, D. R. Morado, G. A. Voth, J. A. G. Briggs, Immature HIV-1 assembles from Gag dimers leaving partial hexamers at lattice edges as potential substrates for proteolytic maturation. Proc Natl Acad Sci U S A 118, (2021).

3. P. Banerjee, V. Monje-Galvan, G. A. Voth, Cooperative Membrane Binding of HIV-1 Matrix Proteins. J Phys Chem B 128, 2595–2606 (2024).

4. P. Banerjee, G. A. Voth, Conformational transitions of the HIV-1 Gag polyprotein upon multimerization and gRNA binding. Biophys J 123, 42–56 (2024).

5. A. Duchon, W. S. Hu, HIV-1 RNA genome packaging: it’s G-rated. mBio, e0086123 (2024).

6. D. Muriaux, J. Mirro, D. Harvin, A. Rein, RNA is a structural element in retrovirus particles. Proc Natl Acad Sci U S A 98, 5246–5251 (2001).

7. J. A. Briggs, J. D. Riches, B. Glass, V. Bartonova, G. Zanetti, H. G. Kräusslich, Structure and assembly of immature HIV. Proc Natl Acad Sci U S A 106, 11090–11095 (2009).

8. K. A. Dilley, O. A. Nikolaitchik, A. Galli, R. C. Burdick, L. Levine, K. Li, A. Rein, V. K. Pathak, W. S. Hu, Interactions between HIV-1 Gag and Viral RNA Genome Enhance Virion Assembly. J Virol 91, (2017).

9. Y. Qiu, P. Banerjee, K. Grabarkewitz, V. H. Wysocki, I. Rouzina, G. A. Voth, K. Musier-Forsyth, Psi RNA-specific Binding Promotes HIV-1 Gag Conformational Change Critical for Immature Viral Particle Assembly. bioRxiv, 2026.2007.2018.739361 (2026).

10. C. Lacouture, B. Carrio, C. Favard, D. Muriaux, HIV-1 assembly - when virology meets biophysics. J Cell Sci 137, (2024).

11. A. J. Pak, J. M. A. Grime, P. Sengupta, A. K. Chen, A. E. P. Durumeric, A. Srivastava, M. Yeager, J. A. G. Briggs, J. Lippincott-Schwartz, G. A. Voth, Immature HIV-1 lattice assembly dynamics are regulated by scaffolding from nucleic acid and the plasma membrane. Proc Natl Acad Sci U S A 114, E10056–E10065 (2017).

12. L. Foret, Shape and energy of a membrane bud induced by protein coats or viral protein assembly. Eur Phys J E Soft Matter 37, 42 (2014).

13. J. H. Hurley, A. K. Cada, Inside job: how the ESCRTs release HIV-1 from infected cells. Biochem Soc Trans 46, 1029–1036 (2018).

14. A. Hudait, J. H. Hurley, G. A. Voth, Dynamics of upstream ESCRT organization at the HIV-1 budding site. Biophys J 122, 2655–2674 (2023).

15. K. Inamdar, F. C. Tsai, R. Dibsy, A. de Poret, J. Manzi, P. Merida, R. Muller, P. Lappalainen, P. Roingeard, J. Mak, P. Bassereau, C. Favard, D. Muriaux, Full assembly of HIV-1 particles requires assistance of the membrane curvature factor IRSp53. Elife 10, (2021).

16. Z. Jarin, F. C. Tsai, A. Davtyan, A. J. Pak, P. Bassereau, G. A. Voth, Unusual Organization of I-BAR Proteins on Tubular and Vesicular Membranes. Biophys J 117, 553–562 (2019).

17. A. Disanza, S. Mantoani, M. Hertzog, S. Gerboth, E. Frittoli, A. Steffen, K. Berhoerster, H. J. Kreienkamp, F. Milanesi, P. P. Di Fiore, A. Ciliberto, T. E. Stradal, G. Scita, Regulation of cell shape by Cdc42 is mediated by the synergic actin-bundling activity of the Eps8-IRSp53 complex. Nat Cell Biol 8, 1337–1347 (2006).

18. J. Saarikangas, H. Zhao, A. Pykäläinen, P. Laurinmäki, P. K. Mattila, P. K. Kinnunen, S. J. Butcher, P. Lappalainen, Molecular mechanisms of membrane deformation by I-BAR domain proteins. Curr Biol 19, 95–107 (2009).

19. F. C. Tsai, J. M. Henderson, Z. Jarin, E. Kremneva, Y. Senju, J. Pernier, O. Mikhajlov, J. Manzi, K. Kogan, C. Le Clainche, G. A. Voth, P. Lappalainen, P. Bassereau, Activated I-BAR IRSp53 clustering controls the formation of VASP-actin-based membrane protrusions. Sci Adv 8, eabp8677 (2022).

20. A. de Poret, R. Dibsy, P. Merida, A. Trausch, K. Inamdar, D. Muriaux, Extracellular vesicles containing the I-BAR protein IRSp53 are released from the cell plasma membrane in an Arp2/3 dependent manner. Biol Cell 114, 259–275 (2022).

21. H. Zhao, A. Pykäläinen, P. Lappalainen, I-BAR domain proteins: linking actin and plasma membrane dynamics. Curr Opin Cell Biol 23, 14–21 (2011).

22. A. M. Chou, K. P. Sem, W. J. Lam, S. Ahmed, C. Y. Lim, Redundant functions of I-BAR family members, IRSp53 and IRTKS, are essential for embryonic development. Sci Rep 7, 40485 (2017).

23. C. Favard, J. Chojnacki, P. Merida, N. Yandrapalli, J. Mak, C. Eggeling, D. Muriaux, HIV-1 Gag specifically restricts PI(4,5)P2 and cholesterol mobility in living cells creating a nanodomain platform for virus assembly. Sci Adv 5, eaaw8651 (2019).

24. N. Yandrapalli, Q. Lubart, H. S. Tanwar, C. Picart, J. Mak, D. Muriaux, C. Favard, Self assembly of HIV-1 Gag protein on lipid membranes generates PI(4,5)P. Sci Rep 6, 39332 (2016).

25. Z. Jarin, A. J. Pak, P. Bassereau, G. A. Voth, Lipid-Composition-Mediated Forces Can Stabilize Tubular Assemblies of I-BAR Proteins. Biophys J 120, 46–54 (2021).

26. Q. Cui, R. Nussinov, Making biomolecular simulations accessible in the post-Nobel Prize era. PLoS Comput Biol 10, e1003786 (2014).

27. R. Evans, M. O’Neill, A. Pritzel, N. Antropova, A. Senior, T. Green, A. Žídek, R. Bates, S. Blackwell, J. Yim, O. Ronneberger, S. Bodenstein, M. Zielinski, A. Bridgland, A. Potapenko, A. Cowie, K. Tunyasuvunakool, R. Jain, E. Clancy, P. Kohli, J. Jumper, D. Hassabis, Protein complex prediction with AlphaFold-Multimer. bioRxiv, 2021.2010.2004.463034 (2022).

28. J. Jumper, R. Evans, A. Pritzel, T. Green, M. Figurnov, O. Ronneberger, K. Tunyasuvunakool, R. Bates, A. Žídek, A. Potapenko, A. Bridgland, C. Meyer, S. A. A. Kohl, A. J. Ballard, A. Cowie, B. Romera-Paredes, S. Nikolov, R. Jain, J. Adler, T. Back, S. Petersen, D. Reiman, E. Clancy, M. Zielinski, M. Steinegger, M. Pacholska, T. Berghammer, S. Bodenstein, D. Silver, O. Vinyals, A. W. Senior, K. Kavukcuoglu, P. Kohli, D. Hassabis, Highly accurate protein structure prediction with AlphaFold. Nature 596, 583–589 (2021).

29. J. B. Ingraham, M. Baranov, Z. Costello, K. W. Barber, W. Wang, A. Ismail, V. Frappier, D. M. Lord, C. Ng-Thow-Hing, E. R. Van Vlack, S. Tie, V. Xue, S. C. Cowles, A. Leung, J. V. Rodrigues, C. L. Morales-Perez, A. M. Ayoub, R. Green, K. Puentes, F. Oplinger, N. V. Panwar, F. Obermeyer, A. R. Root, A. L. Beam, F. J. Poelwijk, G. Grigoryan, Illuminating protein space with a programmable generative model. Nature 623, 1070–1078 (2023).

30. Z. Lin, H. Akin, R. Rao, B. Hie, Z. Zhu, W. Lu, N. Smetanin, R. Verkuil, O. Kabeli, Y. Shmueli, A. Dos Santos Costa, M. Fazel-Zarandi, T. Sercu, S. Candido, A. Rives, Evolutionary-scale prediction of atomic-level protein structure with a language model. Science 379, 1123–1130 (2023).

31. M. J. Abraham, T. Murtola, R. Schulz, S. Pll, J. C. Smith, B. Hess, E. Lindahl, GROMACS: High performance molecular simulations through multi-level parallelism from laptops to supercomputers. SoftwareX 1, 19--25 (2015).

32. J. Jin, A. J. Pak, A. E. P. Durumeric, T. D. Loose, G. A. Voth, Bottom-up Coarse-Graining: Principles and Perspectives. J Chem Theory Comput 18, 5759–5791 (2022).

33. Y. Liu, Z. Yu, R. J. Lindsay, G. Lin, M. Chen, A. Sahoo, S. M. Hanson, ExEnDiff: An Experiment-guided Diffusion model for protein conformational Ensemble generation. bioRxiv, 2024.2010.2004.616517 (2024).

34. J. L. Watson, D. Juergens, N. R. Bennett, B. L. Trippe, J. Yim, H. E. Eisenach, W. Ahern, A. J. Borst, R. J. Ragotte, L. F. Milles, B. I. M. Wicky, N. Hanikel, S. J. Pellock, A. Courbet, W. Sheffler, J. Wang, P. Venkatesh, I. Sappington, S. V. Torres, A. Lauko, V. De Bortoli, E. Mathieu, S. Ovchinnikov, R. Barzilay, T. S. Jaakkola, F. DiMaio, M. Baek, D. Baker, De novo design of protein structure and function with RFdiffusion. Nature 620, 1089–1100 (2023).

35. Y. Xie, L. Winkler, L. Sun, S. Lewis, A. E. Foster, J. Jiménez Luna, T. Hempel, M. Gastegger, Y. Chen, I. Zaporozhets, C. Clementi, C. M. Bishop, F. Noé, Enhanced Diffusion Sampling: Efficient Rare Event Sampling and Free Energy Calculation with Diffusion Models. 10.48550/arXiv.2602.16634 (2026).

36. S. Lewis, T. Hempel, J. Jimenez-Luna, M. Gastegger, Y. Xie, A. Y. K. Foong, V. G. Satorras, O. Abdin, B. S. Veeling, I. Zaporozhets, Y. Chen, S. Yang, A. E. Foster, A. Schneuing, J. Nigam, F. Barbero, V. Stimper, A. Campbell, J. Yim, M. Lienen, Y. Shi, S. Zheng, H. Schulz, U. Munir, R. Sordillo, R. Tomioka, C. Clementi, F. Noe, Scalable emulation of protein equilibrium ensembles with generative deep learning. Science 389, eadv9817 (2025).

37. Z. Yu, Y. Liu, G. Lin, W. Jiang, M. Chen, ESMAdam: a plug-and-play all-purpose protein ensemble generator. bioRxiv, 2025.2001.2019.633818 (2025).

38. J. Ohnuki, K. I. Okazaki, Integration of AlphaFold with Molecular Dynamics for Efficient Conformational Sampling of Transporter Protein NarK. J Phys Chem B 128, 7530–7537 (2024).

39. G. I. Makhatadze, The accuracy of electrostatic interactions captured by AI protein structure prediction models. Proc Natl Acad Sci U S A 123, e2609610123 (2026).

40. G. Scita, S. Confalonieri, P. Lappalainen, S. Suetsugu, IRSp53: crossing the road of membrane and actin dynamics in the formation of membrane protrusions. Trends Cell Biol 18, 52–60 (2008).

41. N. Kurochkina, U. Guha, SH3 domains: modules of protein-protein interactions. Biophys Rev 5, 29–39 (2013).

42. O. Aitio, M. Hellman, A. Kazlauskas, D. F. Vingadassalom, J. M. Leong, K. Saksela, P. Permi, Recognition of tandem PxxP motifs as a unique Src homology 3-binding mode triggers pathogen-driven actin assembly. Proc Natl Acad Sci U S A 107, 21743–21748 (2010).

43. T. R. Gamble, F. F. Vajdos, S. Yoo, D. K. Worthylake, M. Houseweart, W. I. Sundquist, C. P. Hill, Crystal structure of human cyclophilin A bound to the amino-terminal domain of HIV-1 capsid. Cell 87, 1285–1294 (1996).

44. A. Padron, P. Prakash, J. Pandhare, J. Luban, C. Aiken, M. Balasubramaniam, C. Dash, Emerging role of cyclophilin A in HIV-1 infection: from producer cell to the target cell nucleus. J Virol 97, e0073223 (2023).

45. J. C. Ferreon, V. J. Hilser, Thermodynamics of binding to SH3 domains: the energetic impact of polyproline II (PII) helix formation. Biochemistry 43, 7787–7797 (2004).

46. A. Ono, S. D. Ablan, S. J. Lockett, K. Nagashima, E. O. Freed, Phosphatidylinositol (4,5) bisphosphate regulates HIV-1 Gag targeting to the plasma membrane. Proc Natl Acad Sci U S A 101, 14889–14894 (2004).

47. S. A. K. Datta, J. E. Curtis, W. Ratcliff, P. K. Clark, R. M. Crist, J. Lebowitz, S. Krueger, A. Rein, Conformation of the HIV-1 Gag Protein in Solution. Journal of Molecular Biology 365, 812–824 (2007).

48. X. Chen, X. Wang, The HIV-1 gag p6: a promising target for therapeutic intervention. Retrovirology 21, 1 (2024).

49. C. Floderer, J. B. Masson, E. Boilley, S. Georgeault, P. Merida, M. El Beheiry, M. Dahan, P. Roingeard, J. B. Sibarita, C. Favard, D. Muriaux, Single molecule localisation microscopy reveals how HIV-1 Gag proteins sense membrane virus assembly sites in living host CD4 T cells. Sci Rep 8, 16283 (2018).

50. C. Waltmann, A. Shrestha, M. Olvera de la Cruz, Patterning of multicomponent elastic shells by gaussian curvature. Physical Review E 109, 054409 (2024).

51. J. Abramson, J. Adler, J. Dunger, R. Evans, T. Green, A. Pritzel, O. Ronneberger, L. Willmore, A. J. Ballard, J. Bambrick, S. W. Bodenstein, D. A. Evans, C. C. Hung, M. O’Neill, D. Reiman, K. Tunyasuvunakool, Z. Wu, A. Zemgulyte, E. Arvaniti, C. Beattie, O. Bertolli, A. Bridgland, A. Cherepanov, M. Congreve, A. I. Cowen-Rivers, A. Cowie, M. Figurnov, F. B. Fuchs, H. Gladman, R. Jain, Y. A. Khan, C. M. R. Low, K. Perlin, A. Potapenko, P. Savy, S. Singh, A. Stecula, A. Thillaisundaram, C. Tong, S. Yakneen, E. D. Zhong, M. Zielinski, A. Zidek, V. Bapst, P. Kohli, M. Jaderberg, D. Hassabis, J. M. Jumper, Accurate structure prediction of biomolecular interactions with AlphaFold 3. Nature 630, 493–500 (2024).

52. S. William Humphrey and Andrew Dalke and Klaus, VMD: Visual molecular dynamics. Journal of Molecular Graphics 14, 33–38 (1996).

53. L. Deshmukh, C. D. Schwieters, A. Grishaev, R. Ghirlando, J. L. Baber, G. M. Clore, Structure and dynamics of full-length HIV-1 capsid protein in solution. J Am Chem Soc 135, 16133–16147 (2013).

54. C. Waltmann, Y. Wang, C. Yang, S. Kim, G. A. Voth, MSBack: Multiscale Backmapping of Highly Coarse-Grained Proteins Using Constrained Diffusion. J Chem Theory Comput 21, 6184–6193 (2025).

55. S. Kim, Backmapping with Mapping and Isomeric Information. J Phys Chem B 127, 10488–10497 (2023).

56. M. Jones, S. Khanna, A. Ferguson, Flowback: A Flow-matching Approach for the Generative Backmapping of Macromolecules. (2024).

57. W. L. Jorgensen, J. Chandrasekhar, J. D. Madura, R. W. Impey, M. L. Klein, Comparison of Simple Potential Functions for Simulating Liquid Water. Journal of Chemical Physics 79, 926–935 (1983).

58. J. Huang, S. Rauscher, G. Nawrocki, T. Ran, M. Feig, B. L. de Groot, H. Grubmuller, A. D. MacKerell, Jr., CHARMM36m: an improved force field for folded and intrinsically disordered proteins. Nat Methods 14, 71–73 (2017).

