## Supplemental Figures 1-5 for "HIV-1 Assembly is Assisted by IRSp53 Through an Interaction that Extends Gag Conformation"

#### Additional Figures

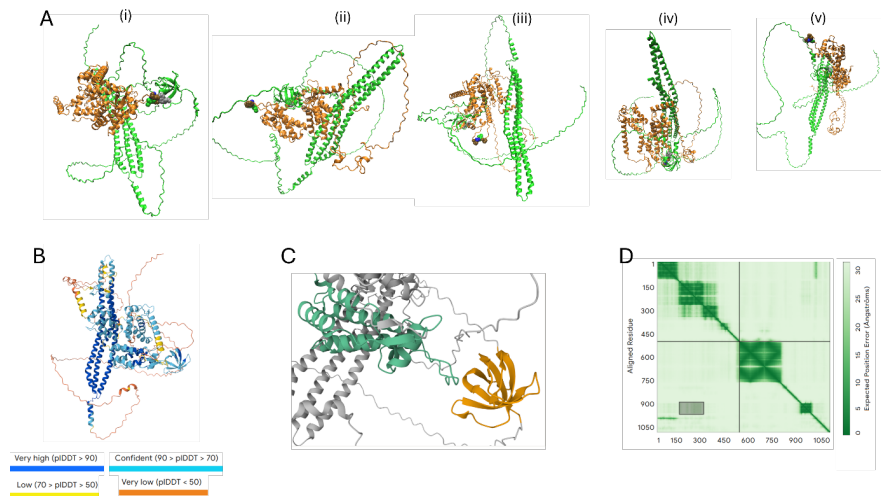

**Figure S1.** Structural Analysis of AlphaFold predictions. (A) The full-length structure of Gag (Orange) and IRSp53(green) for the 5 most confident AF predictions. The PIAP domain on Gag and the TRP which was predicted to insert into it are shown in van der Waals representation. (i) The structure that was chosen as an initial guess shows the TRP inserted into the PIAP. For (ii-v) these two motifs are not close to one another. (B) The first structure is shown with the predicted local distance difference test (pLDDT) scores. (C) Shows a zoomed in image of the gag CATND (green) with the IRSP53 SH3 domain (orange). (D) The expected position error scores for all pairs of residues (numbered Gag then IRSP53). There is some confidence in the distances in the SH3-CANTD highlighted region (also look at the upper triangle) but not too much.

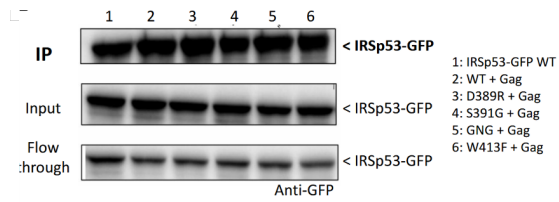

**Figure S2:** Calibration of the assay from Figure 3G

Commented [CW1]: Need Delphine here

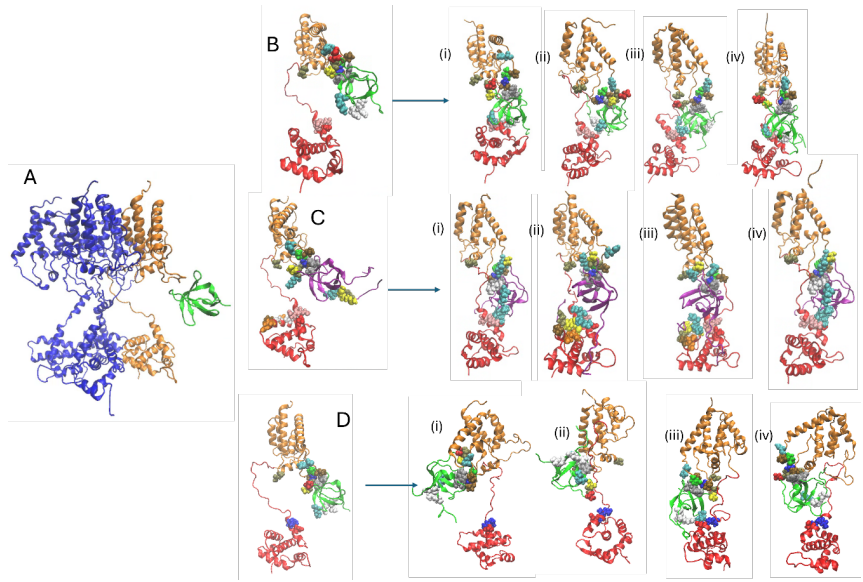

**Figure S3.** Simulations with Gag MACANTD constrained to simulate the assembled state. (A) Snapshot of the IRSP53 domain bound to one gag (orange) of the gag 4-mer (the other 3 gags are blue). The SH3 domain is bound to the CANTD as it is in Figure 2 in the main text. (B) The initial simulation state which removes the 3 blue gags and replaces them with harmonic restraints that simulate the gag-gag contacts. Snapshots show binding of gag to the MA domain of gag as well as the CANTD for all 4 independent replicas (i-iv). (C) The same initial state but the IRSP53 SH3 is replaced with the IRTKS SH3 domain. In (i-iv) binding of the IRTKS SH3 domain the MA and CANTD is also shown for all 4 independent replicas with similar contacts despite sequence differences between IRTKS and IRSP53. Here, we cannot establish a difference between IRSP53 and IRTKS. (D) The same initial state but the Gag is replaced with 3E3A Gag explained in the main text. We ran this mutant since the E residues mutated were important in binding the IRSp53 SH3 domain with WT Gag. In (i-iv), only (iii) shows binding of the IRTKS SH3 domain to the MA domain. All IRSp53 SH3 domains remain bound to the CANTD for the 4 independent replicas for the full 500 ns simulation.

#### SI Figure 4. Sequences

##### Gag Sequence Used for Alphafold Predictions – Key Residues Highlighted

MGARASVLSGGELDKWEKIRLRPGGKKQYKLKHIVWASRELERFAVNPGLLETSEGCRQILGQ
LQPSLQTGSEELRSLYNTIAVLYCVHQRIDVKDTKEALDKIEEEQNKS**KKKAQQ**AAADTGNNSQ
VSQ

NYPIVQNLQGQMVHQAISPRTLNAWVKVVEEKAFSPEVIPMFSALSEGATPQDLNTMLNTVGG
HQAAMQMLKETINEEAAEWDRLHPV**HAGPIAP**GQMREPRGSDIAGTTSTLQEQIGWMTH**N**PPI
PVGEIYKRWIILGLNKIVRMYSPTSILDIRQGPKEPFRDYVDRFYKTLRAEQASQEVKNWMTETL
LVQNANPDCKTILKALGPGATLEEMMTACQGVGGPGHKARVLAEAMSQVTNPATIMIQKGNFR
NQRKTVKCFNCGKEGHIAKNCRAPRKKGCWKCGKEGHQMKDCTERQANFLGKIWPSHKGRP
GNFLQSRPEPTAPPEESFRFGEETTTPSQKQEPIDKELYPLASLRSLFGSDPSSQ

##### Gag sequence used for MACANTD/CANTD simulations – Difference with AF Sequence Highlighted

**Y**ARASVLSGGELDKWEKIRLRPGGKKQYKLKHIVWASRELERFAVNPGLLETSEGCRQILGQ
LQPSLQTGSEELRSLYNTIAVLYCVHQRIDVKDTKEALDKIEEEQNKS**KKKAQQ**AAADTG**HSN**Q
VSQ

NYPIVQN**I**QQGMVHQAIPTLNNAWVKVVEEKAFSPEVPMFSALESEGATPQDLNMLNTVGG
HQAAMQMLKETINEEAAEWDRLHPV**HAGPIAP**GQMREPRGSDIAGTTSTLQEQIGWMTHNPPI
PVGEIYKRWIILGLNKIVRMYSPTS

##### Sequence of IRSP53 Used for Alphafold and MD Simulation (Uniprot Q9UQB8 BAIP2\_HUMAN)

MSLSRSEEMHRLTENVYKTIMEQFNPSLRNFIAMGKNYEKALAGVTYAAKGYFDALVKMGELA
SESQGSKELGDVLFQMAEV
HRQIQNQLEEMLKSFHNELLTQLEQKVELDSRYLSAALKKYQTEQRSKGDALDKCQAELKKLR
KKSQGSKNPQKYSDKELQYIDAISNKQGELENYVSDGYKTALTEERRRFCFLVEKQCAVAKNS
AAYHSKGKELLAQKLPLWQQACADPSKIPERAVQLMQQVASNGATLPSALSASKSNLVISDPIP
GAKPLPVPPELAPFVGRMSAQESTPIMNGVTGPDGEDYSPWADRKAAQPKSLSPPQSQSKLS
DSYSNTLPVRKSVTPKNSYATTENKT

LPRSSSM (Added to check linker effect in some replicas)

##### **Begin SH3**

AAGLERN (Not in CANTD only simulation)

GRMRVKAIFSHAAG**DNS**TLLSFKEGDLITLLVPEARDGWHYGESEKTKMRGWFPFSYTRVLD
SDG

##### **End SH3**

SDRLHMSLQQGKSSSTGNLLDKDDLAIPPPDYGAASRAFPAQTASGFKQRPYSVAVPAFSQGL
DDYGARSMSRNPFAHVQLKPTVTNDRCDLSAQGPEGREHGDGSARTLAGR

Sequence of IRTKS SH3 For MD Simulations (from Uniprot Q9UHR4 BI2L1 HUMAN and PDB 2KXC)

VSV (added to check linker effect in some replicas)

**Begin SH3**

ATGLN (Used with MACANTD GAG) – GSH (Used with CANTD only, artifact from PDB 2KXC)

MMKKQKVKTIFPHTAGSNKTLLSFAQGDVITLLIPEEKDGWLYGEHDVSKARGWFPSSYTKLLE
ENE

**End SH3**

TEAVTV (added to check linker effect in some replicas)

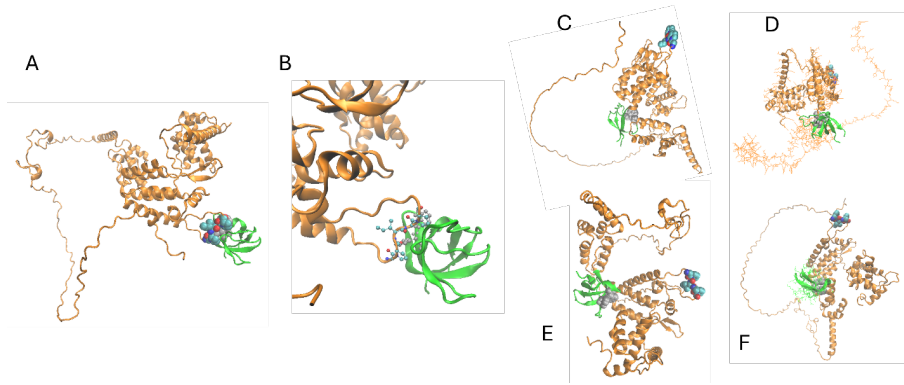

*Figure S5. AF2 predictions for full-length Gag with IRSp53 SH3. (A) The most confident prediction of the full-length structure of Gag (orange) and IRSp53 SH3 (green). The PIAP domain on Gag and the TRP which was predicted to insert into it are shown in van der Waals representation. (B) The same structure enlarged to see the interaction domain which is nearly identical to the AF3 prediction with both full-length sequences. (C-F) The lower ranked structures do not show reasonable interactions. There are structural clashes and the SH3 domain is positioned between the NTD and CTD of CA, which does not fit the assembled Gag lattice.*

**Supplemental Methods**

Restraining Gag in the multimerized state was done to mimic the effect of the rest of the Gag lattice on the gag protein conformation. A harmonic potential for each cardinal direction was placed on the absolute position of the alpha carbon of a series of amino acids with a 50 kJ/mol/nm<sup>2</sup> bond constant. Those residues were 1-10, 44-46, 69-71, 149, 153, 157, 161, 168, 169, 173, 207, 267, 270, and 274. Additionally, in two of the replicas with IRSP53 and three of the replicas with IRTKS the position of residues 26, 27, and 29 were also restrained to model the interaction between these charged residues (sometimes called the highly basic region, HBR) and the membrane as noted in the text we did not see any effects on SH3 binding based on how the HBR was handled.
